# Long-term citizen science data reveals environmental correlates of tropical tree flowering at the regional scale

**DOI:** 10.1101/2023.03.24.533907

**Authors:** Krishna Anujan, Jacob Mardian, Carina Luo, Ramraj Rajakumar, Hana Tasic, Nadia Akseer, Geetha Ramaswami

## Abstract

Tropical tree reproductive phenology is sensitive to changing climate, but inter-individual and interannual variability at the regional scale is poorly understood. While large-scale and long-term datasets of environmental variables are available, reproductive phenology needs to be measured in-site, limiting the spatiotemporal scales of the data. We leveraged a unique dataset assembled by SeasonWatch, a citizen-science phenology monitoring programme in India to assess the environmental correlates of flowering in three ubiquitous and economically important tree species - jackfruit, mango and tamarind - in the south-western Indian state of Kerala. We explored (i) seasonal patterns in the flowering status of trees (ii) environmental correlates of flowering onset considering only trees with consecutive observations and (iii) spatiotemporal patterns in these environmental correlates to aid future hypotheses for changing phenology patterns. We used 165006 phenology observations spread over 19591 individual trees over 9 years. We first used bootstrapped circular statistics that accounts for observation biases in time to examine consistency in seasonality of flowering status over the whole season, and flowering onset across years and trees. Similar to results from cohort-based tree monitoring, we demonstrate seasonality in flowering status and onset across species, but also report large interannual and inter-individual variability. We then used used generalized linear mixed models with remotely sensed observations (ERA5-LAND) to show that some of the interannual variation in flowering onset across individuals was associated with environmental variables. Soil moisture, minimum temperature and solar radiation had significant associations with the onset of flowering but these effects were heterogeneous across species and habitats across Kerala. Our results become increasingly important in the face of large spatiotemporal change in the climate of this landscape and other tropical regions. We demonstrate the potential and limitations of citizen- science observations in making and testing predictions at scale for predictive climate science in tropical landscapes.

**Open Research Statement:** Data are not yet provided. Environmental data used in this manuscript are from publicly available sources. All SeasonWatch data, the phenology data used in the manuscript, is licensed under the open-access creative commons license CC BY 4.0: https://creativecommons.org/licenses/by/4.0/ and is available upon request. Processed data and code used in this manuscript will be made publicly accessible through Zenodo upon acceptance of the manuscript. This manuscript does not use any novel code.

## Introduction

Tree reproductive phenology is altered by a changing climate, affecting species interactions and ecosystem functions and services (Cleland et al. 2007, Mora et al. 2015). Flowering in trees is triggered by environmental cues like rainfall events, dry spells, and photoperiod, and is therefore sensitive to a changing environment (Chapman et al. 2005, Chang-Yang et al. 2016, Pau et al. 2018). Changes in onset, persistence, and quantity of reproductive phenophases can alter flower and fruit availabilty in the landscape, affecting interactions with other species like pollinators, frugivores, and seed dispersers (Visser and Both 2005, CaraDonna et al. 2014, Morellato et al. 2016). Consequently, for wild populations of trees, changes in phenology can influence regeneration dynamics and population persistence. Changing phenology can also have socio- cultural impacts, through altering food production and creating mismatches in cultural understanding of tree behaviours (Mora et al. 2015, Morellato et al. 2016). While landscape-level phenological shifts in temperate regions have been studied over decades, large gaps remain in ecological documentation and predictions for shifts in the reproductive phenology of tropical trees.

In tropical landscapes where trees grow and reproduce year-round, environmental cues for phenological change can be spatially and temporally diffuse (Parmesan 2007, Chen et al. 2018, Abernethy et al. 2018). There is a growing understanding of the influence of environmental factors on leaf phenology of tropical species, but information on reproductive phenology is lacking (Chaturvedi et al. 2021, Devi et al. 2023). Cohort monitoring of trees has shown that temperature, precipitation, and solar radiation (e.g. cold days, dry spells, total accumulated rainfall and soil moisture) affect the onset and persistence of flowering phenology in tropical trees (Singh and Singh 1992, Brearley et al. 2007, Pau et al. 2013, Diez et al. 2014, Wright and Calderón 2018). Interannual rainfall variability - high rainfall years and drought years including cycles like the El Niño Southern Oscillation and the Indian Ocean Dipole - can result in interannual variability in tree phenophase onset and intensity (Medway 1972, Zimmerman et al. 2007, Chang-Yang et al. 2016, Chapman et al. 2018, Dunham et al. 2018, Pau et al. 2018).

Furthermore, climate change has resulted in altered rainfall and temperature regimes as well as extreme events in tropical environments, increasing inter-annual variability (Mora et al. 2013). To predict future phenological shifts in the tropics, an important first step is to understand the extent of environmental controls on tropical tree reproductive phenology.

Environmental cues could affect phenology with a lag time, interact with habitat factors and affect individuals differently, necessitating large-scale and long-term data. In tropical species, at least six years of data is necessary to accurately detect phenology in regularly flowering species, with much more for noisy species (Bush et al. 2017). Species across habitats may have different responses to the same environmental cues, with differences in timing due to urbanisation (Neil and Wu 2006), variation in synchrony with latitude (Borchert et al. 2004) or with distance to coast (Vanderplank and Ezcurra 2016). Tropical tree phenology also exhibits large variation among individuals, necessitating large spatial samples besides long-term data for a process-based understanding(Satake et al. 2013, CaraDonna et al. 2014, Li et al. 2021). A recent review of reproductive phenology studies from the Neotropics showed that across 218 datasets, the majority of datasets lasted 2 years or less, and only 10.4% monitored > 15 individuals per species (Mendoza et al. 2017). Across the tropics, phenology of ubiquitous species have rarely been recorded systematically, hindering the potential to make large-scale predictions of climate impacts of tree-associated ecosystem services (Piao et al. 2019, McDonough MacKenzie et al. 2020). In the absence of robust datasets, potentially high intraspecific variation in reproductive phenology makes it difficult to parametrise models that span spatial scales of environmental change.

Citizen science allows access to longitudinal data across large spatial scales that is difficult to achieve through other sampling techniques. Citizen science data has led to novel understanding of phenological patterns in other taxa - observations of migratory bird populations across North America have been useful in documenting phenological shifts, spatial differences in sensitivities to changing climate, as well as disentangling drivers of change (Hurlbert and Liang 2012, Kelly et al. 2016, Haas et al. 2022). Other efforts, like the USA National Phenology Network, have helped quantify advancing spring behaviours of plant species in North America (Schwartz et al. 2012, Brunsdon and Comber 2012). SeasonWatch, a citizen science phenology monitoring programme has been collecting data on several phenophases for common, widespread tree species across India (Ramaswami et al. 2019, 2021). Each week, observers record flowering, along with other phenophases, on specific trees consistently using a uniform protocol that records both presence and intensity of a phenophase (Fig 1a). This dataset, with substantial contributions since 2014, provides a unique opportunity to build baselines for phenology in the Indian subcontinent. In the past, some analyses of these data have revealed patterns of change in at least one species - Cassia fistula - over the time period of observation (Ramaswami and Quader 2018, Ramaswami et al. 2021). These common species, spread across ecoregions and habitats, can add to phenological understanding at large scales, complementing focussed, localised studies.

**Figure 1:**
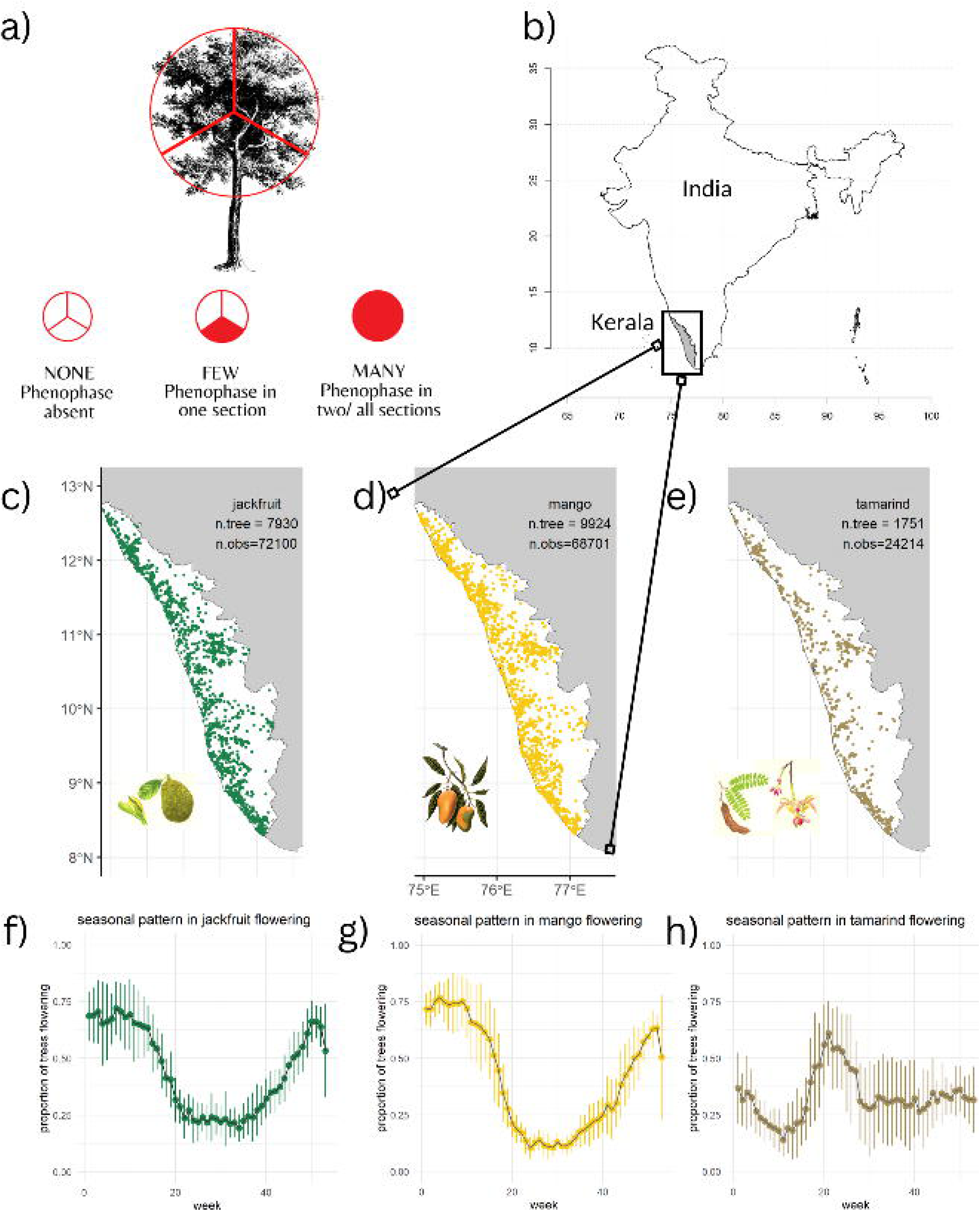
SeasonWatch protocol, species and map of observations. a) SeasonWatch observation protocol for tree phenology - phenophase quantities ‘none’, ‘few’, ‘many’ roughly correspond to the presence of a phenophase in 0, 30%, and >30% of the tree canopy b) Outline of India with the state of Kerala marked in grey. c), d) and e) - map of observations of jackfruit, mango and tamarind trees across Kerala, considered in this study along with illustrations of flowering and fruiting phenophases in jackfruit, mango and tamarind respectively. Panels f), g) and h) show seasonal patterns in flowering of the three species in Kerala from 2014-2022. Dots represent mean proportion of trees flowering in that week and error bars represent standard deviation across the years of study. Image credits: a) Sayee Girdhari for SeasonWatch; illustration in c) and e) Neelam Modi for SeasonWatch; illustration in d) Mango (Mangifera indica L.): fruiting branch with numbered sections of flower and seed. Chromolithograph by P. Depannemaeker, c.1885, after B. Hoola van Nooten. Wellcome Collection. Source: Wellcome Collection Licence: Public Domain Mark.

Here, we investigated the environmental correlates of flowering in common tree species in India at the individual tree scale using SeasonWatch data. We used 165006 phenology observations across 19591 individual trees over 9 years from tree species in the southwestern Indian state of Kerala, and correlated these with remotely-sensed environmental variables using machine learning and generalised linear mixed model approaches. We asked (i) what are the seasonal patterns in flowering onset and flowering status of common trees across Kerala and what is the extent of interannual and interindividual variation? (ii) What are the environmental correlates associated with tree-level flowering onset? (iv) How are relevant environmental cues changing spatiotemporally across the landscape? We focussed on flowering of three species with the greatest number of individuals and observations across Kerala - jackfruit (*Artocarpus heterophyllus*), mango (*Mangifera indica*) and tamarind (*Tamarindus indica*). We expected (i) interannual and interindividual variation in flowering status and onset to varying degrees across the three species, (ii) based on research using cohort sampling of trees, we expected temperature and precipitation patterns to influence the onset of flowering. (iii) However, given the large spatial extent of our dataset, we also expected that tree flowering responses to environmental correlates would vary with elevation, aspect, and degree of urbanisation. (iv) Finally, through exploratory analysis, we expected to observe and quantify spatiotemporal variation in environmental correlates of flowering phenology in Kerala.

## Methods

### Study Area, Species and SeasonWatch Data

We analyzed phenology and environmental data from the south-western Indian coastal state of Kerala. Bounded between 8° 18’ N, 12° 48’ N, 74° 52’ E and 72° 22’ E, Kerala lies along the Arabian Sea with a tropical monsoon climate. The total population of the state is over 3 million, spread over an area of 38,863 km^2^. The state receives an average annual rainfall of over 3000mm via two major branches of the Indian monsoon system - the southwest (June-September) and northeast (October-December) monsoon. Citizen scientists from the state have contributed substantially to phenology data; across all SeasonWatch data (SeasonWatch Citizen Scientist Network, 2022, Ramaswami (2022)), 263,288 observations across species and categories, and including 93.23% of observations of top three species are from Kerala. These data span a large range of the existing environmental variability in the state in terms of rainfall, temperature, elevation.

We analyzed flowering phenology in three frequently observed species in the SeasonWatch dataset - *Artocarpus heterophyllus* (henceforth “jackfruit”), *Mangifera indica* (henceforth “mango”) and *Tamarindus indica* (henceforth “tamarind”) from January 2014 - July 2022. All three tree species are grown across the state in orchards, homes, and also occur wild. As part of SeasonWatch, observers register one or more “regular” trees from a list of suggested species on a mobile phone application or web browser and record phenology on a weekly basis. The project also allows for “casual” or one-time observations of trees (outside an observer’s “regular” list), often during coordinated events, expanding the spatial coverage of the data. All observations to SeasonWatch comprise estimates of quantity (“none”, “few”, “many”) of leaf (young, mature, dry), flower (bud, open flower) and fruit (unripe, ripe) phenophases of a tree. For each of the 7 phenophases, the standard protocol for reporting involves dividing the canopy of a tree into thirds, reporting as ‘none’ if it is not observed, as ‘few’ if a phenophase is observed only in 1/3rd of the canopy, and as ‘many’ if a phenophase is observed on more than 1/3rd of the canopy (Fig 1a). SeasonWatch data is contributed by voluntary participants; the programme recruits citizen scientists by first identifying regional partner organisations, training the trainers in these organisations who then recruit and train participants (e.g. teachers, students and community members), thus amplifying the network reach. The project minimises inter-observer biases and errors in observation through workshops and training as well as one-on-one follow-up and troubleshooting. Pertinently, the flowering phases of the common trees chosen for analysis are distinctly observable with the naked eye, and familiar to participants, lending confidence to community data. The data distribution across species and years is provided in Table 1, Appendix S1: Fig S1 and Appendix S1: Fig S2.

**Table 1:** Data distributions. Description of the SeasonWatch dataset for jackfruit, mango and tamarind in Kerala from January 2014 to July 2022

|  | Artocarpus heterophyllus | Mangifera indica | Tamarindus indica |
| --- | --- | --- | --- |
| Total number of observations | 72,097 | 68,696 | 24,213 |
| Total number of trees | 7,927 | 9,919 | 1,750 |
| Number of trees with single observations | 6,270 | 8,128 | 1,277 |
| Number of observations with no flowering at t-1 | 25,695 | 26,212 | 12,426 |
| Number of trees in observations with no flowering at t-1 | 1,028 | 1,105 | 320 |

### Data cleaning and processing

We cleaned and processed SeasonWatch data for jackfruit, mango and tamarind from 2014-2022 to create two different data subsets - flowering status and flowering onset. To maximise inclusion of data points, we included all “regular” and “causal” observations that we were able to reliably assign and validate locations (see Appendix S1: Section S2). We recoded the none/few/many observations of bud and flowers into presence/absence of flowering by coding an observation as “1” if it was recorded as “few” or “many” for buds or flowers, and “0” otherwise. We chose to combine the bud and the flowering phenophases to minimise observation errors in these species where flowers are small and/or greenish and can be difficult to distinguish from buds, especially in taller canopies. Our choice of binary variable also reduces potential user subjectivity in the categorising of “few” and “many”.

#### Flowering status

Flowering status included all observations across trees and years in the dataset where presence or absence of flowering (bud or flower) was recorded. We assembled a final flowering status dataset of 165006 observations across 19596 trees; 7927 trees of jackfruit, 9919 of mango and 1750 of tamarind across the state of Kerala (Fig 1, Appendix S1: Table S1). The number of observations varied by week and year, with numbers generally increasing in the later part of the study period because of increased participation in the programme. (Appendix S1: Fig S1) Most trees in this dataset were observed few times, but several trees had long, uninterrupted flowering status timeseries for one or multiple years (Appendix S1: Fig S4).

#### Flowering onset

The onset of flowering is defined as the first week in a year when a tree switches in status from non-flowering to flowering. To estimate true flowering onset at the tree level, we filtered the flowering status dataset to exclude all observations for which there was no flowering status observation in the preceding week, thus minimising observation bias. Since we were interested in the drivers of the switch from non-flowering to flowering, we further filtered this dataset to only include observations when the tree was observed to be non-flowering in the previous week. For trees that were observed to have multiple onsets in a year (starting from June, the beginning of the wet season), we retained only the first onset observation. This reduced the dataset to 62721 observations with an immediate predecessor, across 2442 trees, out of which, 2638 were true onset observations (0, 1) and the remaining were status quo observations (0, 0) (Table 1, Appendix S1: Fig S2). By filtering to only observations with a non-flowering predecessor, the flowering onset dataset has minimal temporal autocorrelation among observations, allowing the application of mixed modelling frameworks.

### Environmental datasets

We derived seven potential spatiotemporal predictors of flowering from remotely sensed data - three related to temperature (minimum, maximum and mean), three to precipitation (total precipitation, soil moisture and the number of consecutive dry days), and solar radiation. We used daily measurements of precipitation, temperature, solar radiation and soil moisture for the entire soil column from the ERA5-Land dataset at the 11.1 km spatial resolution and summarised these to the fortnight (14 days) preceding each observation using Google Earth Engine (Table 2). Monthly and quarterly summaries were highly correlated to the fortnightly value (Appendix S1: Fig S3) and hence we used this finest temporal resolution (fortnight) for the analysis. Along with spatiotemporal predictors, we calculated spatial factors - elevation, aspect and degree of urbanisation - at each tree location that might potentially act as modifiers for the temporal predictors (Table 2). We used a Digital Elevation Model created from the Shuttle Radio Topography Mission (SRTM) at the 30 m resolution to calculate elevation, slope and aspect at the tree location. Further, we used the NASA nightlights layer from 2016 at the 500 m resolution as a proxy for the degree of urbanisation (Sutton et al. 2001).

**Table 2:**
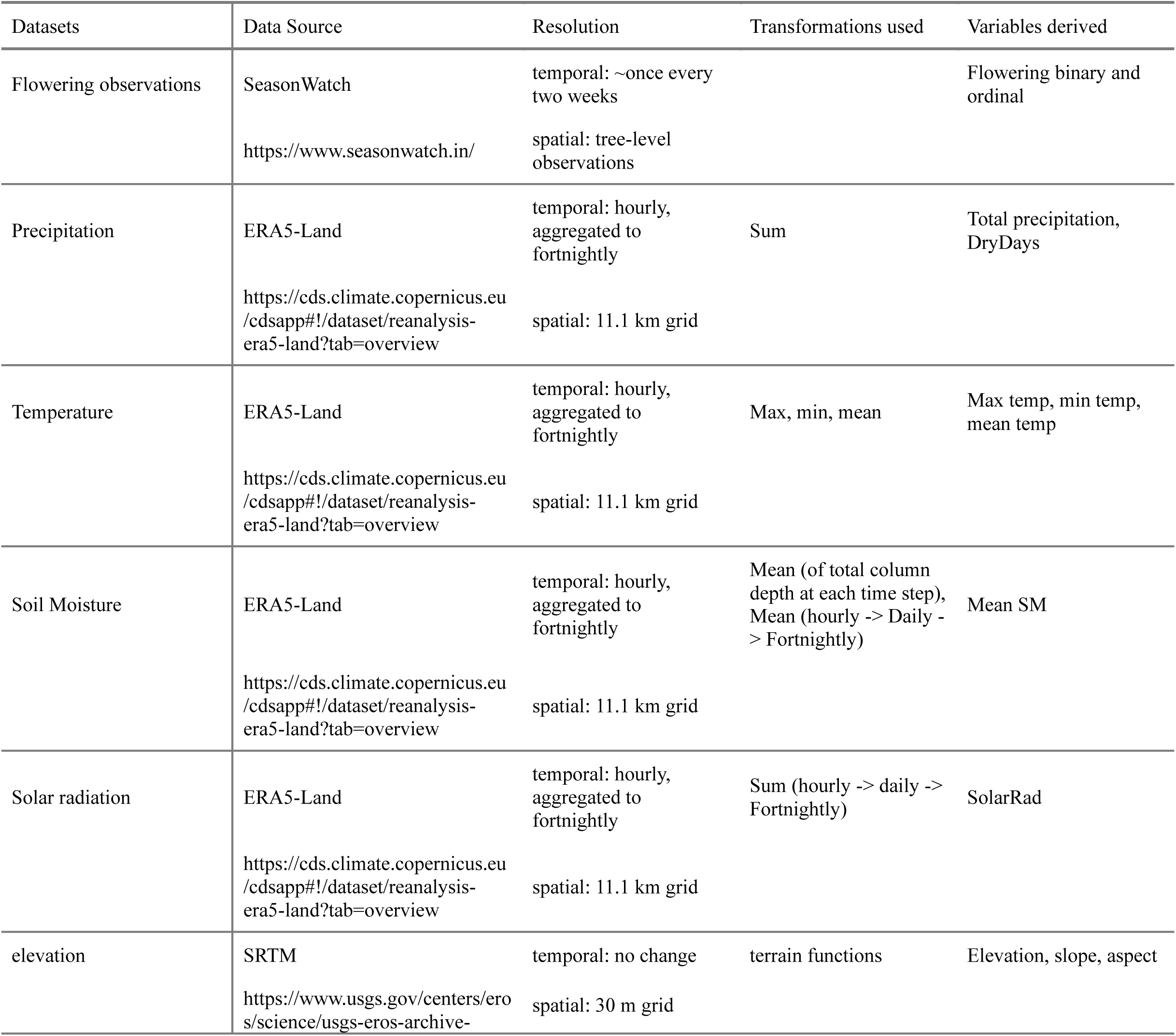
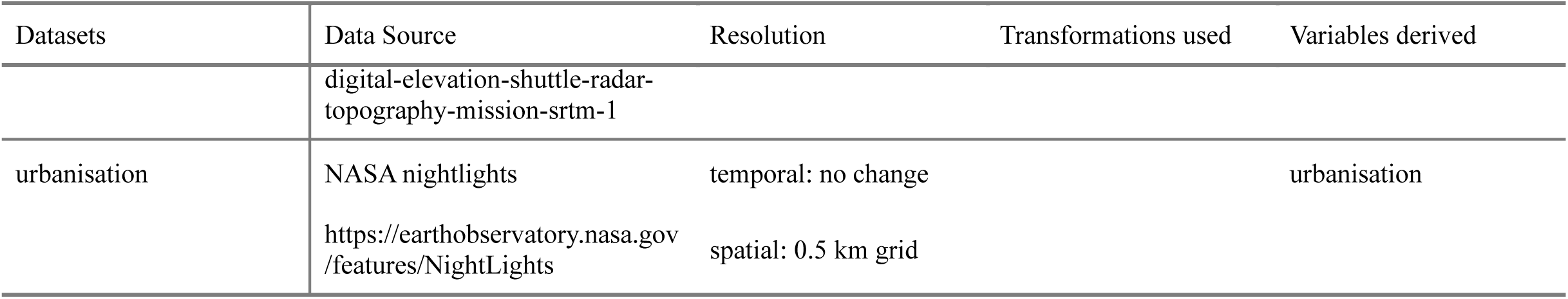
Environmental data. Variables, data sources and transformations

### Data analysis

We analysed seasonality in flowering status and flowering onset at the population level using circular statistics and then modelled predictors of individual-level flowering onset across years of observations using generalised linear mixed models. We modelled flowering seasonality and the ecological predictors of flowering onset separately for each of the three species, as these species are physiologically and phylogenetically distinct (belonging to families - Moraceae, Anacardiaceae, and Fabaceae) and may therefore show distinct flowering seasonality and heterogeneous responses to the same environmental cues. Finally we examined spatiotemporal patterns in environmental predictors across Kerala using hotspot analyses. Analyses were performed using R version 4.1.3 and ArcGIS Pro version 3.0.2 (R Core Team 2022).

### Circular statistics to estimate population flowering seasonality and onset seasonality

We used a newly developed bootstrap method to assess seasonality in flowering in the three species across the state of Kerala (Willig et al. 2024). Seasonality in flowering is best described using circular statistics like circular mean and the Rayleigh statistic which describe the clustering of the data in circular axes (here, week of year) (Morellato et al. 2010). Unlike long-term field monitoring datasets which have fixed cohorts that are repeatedly monitored during the entire monitoring period, citizen science datasets like SeasonWatch can have a) different start and end dates for different individuals b) different timeseries lengths and c) larger proportion of missingness, despite greater spatial spread. This unique data structure leads to non-uniform data distribution across the temporal extent and makes this analogous to capture sampling of random individuals in a population at a fixed frequency, violating the uniformity assumptions of traditional circular statistics (Willig et al. 2024). Using a bootstrapping method, we calculated the circular mean of flowering status and flowering onset for each year of study. We sampled observations with replacement for each week of study to a constant N, equal to the average weekly observation for that species, making the sample distribution uniform across the year.

With this uniform sample, we calculated circular mean, Rayleigh statistic (a measure of temporal clustering) and the p-value of the statistic. We repeated the sampling for 1000 iterations and calculated the bootstrap mean and deviation. To examine interannual consistency in flowering onset across individuals, we calculated mean week of flowering onset and standard deviation for each tree across years all years of onset observations.

### Generalised linear mixed models for flowering onset at the individual level

We examined the effects of environmental variables on the onset of flowering in the three species using generalised linear mixed effects models for interactions between variables and variable collinearity. For each of the three species modelled, the time-varying predictors were identified by picking the most influential variables from random forest (machine learning) models using all environmental variables as predictors (Appendix S1: Section S4). For each species, we picked most important variable in the temperature group (among minimum, maximum and mean), the most important variable in the precipitation group (among total precipitation, number of dry days and soil moisture), and solar radiation. For all species, the most influential temperature variable was minimum temperature and precipitation variable was soil moisture. This grouping allowed us to test the effects of temperature, precipitation and solar radiation, while avoiding including highly correlated variables that can violate mixed modelling assumptions.

We accounted for interactions between time-varying and static predictors, excluding known, correlated variable combinations. Specifically, we excluded interactions between elevation and minimum temperature, aspect and precipitation (the Western Ghats range in Kerala affects rainfall patterns), degree of urbanisation and minimum temperature (known heat island effects in urban areas), degree of urbanisation and soil moisture (potentially associated with heat island effects). Further, to account for data structure in space and time, we included random effects of tree and the spatial grid in which it was located.

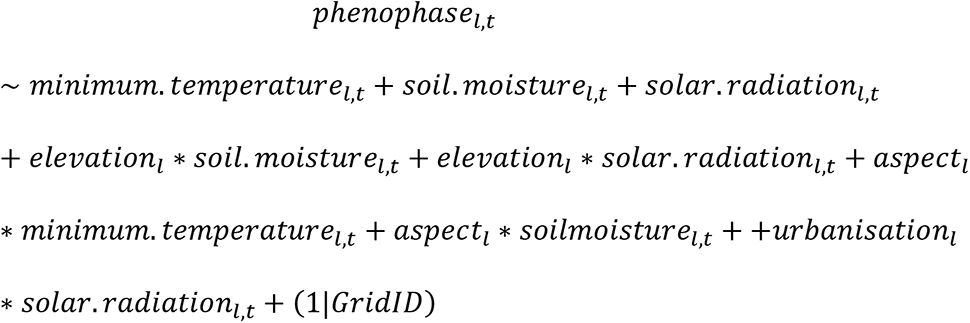

Where minimum temperature, soil moisture and solar radiation are measured at the location 𝑙 for the fortnight before the observation 𝑡, elevation, aspect and urbanisation are spatial predictors tied to the tree location 𝑙. The model includes two random effects on the intercept; 𝑇𝑟𝑒𝑒𝐼𝐷 to account for repeated measures on the same tree and 𝐺𝑟𝑖𝑑𝐼𝐷 to account for spatial autocorrelation.

We modelled the onset of the phenophase using a generalized linear mixed model with a binomial family of errors using the *package* lme4 (Bates et al. 2015). For all models, we mean-centred all predictor variables at 0 and scaled it to a distribution with standard deviation 1 for ease of comparing the effect sizes. We tested the fit of these models using conditional and marginal R^2^ using packages *performance* and *sjPlot* (Lüdecke et al. 2021, Lüdecke 2023). We used a cross-validation technique, leave-one-out cross-validation (LOOCV), to test the predictive power of our model for “out-of-bag” observations. In this method, we ran each model 9 times, systematically leaving one year of observations out each time. We then used the model coefficients to predict these “new” observations and tested their accuracy, and then tested for reliability of predicted values using F1 scores. F1 scores are good indicators of the distribution of true and false positives and negatives by taking the harmonic mean of precision and recall (Hossin and Sulaiman 2015, Grandini et al. 2020).

### Hotspot analysis

We analysed spatiotemporal patterns of environmental variables that affect flowering onset in Kerala, with an intention to identify directional changes or spatial shifts. Hotspot analysis allows the identification of such spatiotemporal patterns using the entire timeseries of environmental data across the years measured. A hot spot/cold spot is defined as a location which has high (or low) value for an environmental variable, like temperature, precipitation or soil moisture, and is surrounded by other areas with similar values; a spatial cluster. Using fortnightly aggregated environmental variables from 2014-2022, we ran emerging hotspot analysis in ArcGIS Pro for minimum temperature, mean soil moisture and total solar radiation, the three variables used for the GLMM analysis, across Kerala. We used a combination of two statistical measures 1) the Getis-Ord Gi* statistic to identify hot spots and cold spots of climate measurements for each time step (i.e., fortnightly), and 2) the Mann Kendall trend test to examine how hot spots and cold spots have evolved over time (Mann 1945, Sen 1968, Kendall et al. 1987, Getis and Ord 1992). For further details on methods, refer Appendix S1: Section S7.

## Results

### Seasonality in flowering across Kerala

Jackfruit, mango and tamarind trees exhibit differing patterns of flowering seasonality and substantial variation among years in mean flowering and flowering onset in the state of Kerala. Across 9 years of data (2014 to 2022), bootstrapped circular mean of flowering status showed that mean flowering week was week 4.88 for jackfruit, 4.68 mango and 25.4 for tamarind.

Temporal clustering in flowering across all years was low, but jackfruit and mango showed higher clustering in flowering across the population than tamarind; bootstrapped Rayleigh statistic, *z* = 0.29 (95% CI = 0.29, 0.3) for jackfruit, 0.41 (95% CI = 0.41, 0.42) for mango and 0.1(95% CI = 0.09, 0.11) for tamarind. Correspondingly, mean week of flowering onset was week 0.57 for jackfruit, 0.4 for mango and 19.45 for tamarind. Patterns of temporal clustering in flowering onset mirrored flowering status; Rayleigh statistic, *z* = 0.5 (95% CI = 0.48, 0.52) for jackfruit, 0.52 (95% CI = 0.5, 0.54) for mango and 0.19 (95% CI = 0.14, 0.24) for tamarind.

Across all three species, trees were observed to be flowering at all times of the year; no week had a zero proportion of trees flowering (Fig 1).

Across 8 full years of observations (excluding the first half of 2014), mean week of flowering status and flowering onset showed interannual variation, but there was low variation in temporal clustering of flowering (Fig 2). Mean flowering week ranged from week 2.73 in 2019 to 7.8 in 2016 for jackfruit, 2.39 in 2021 to 7.35 in 2016 for mango and −25.15 in 2019 to 25.47 in 2015 for tamarind (Fig 2a). Clustering in flowering status was relatively similar across the years (Fig 2a). For jackfruit, bootstrapped Rayleigh statistic of flowering status ranged from *z* = 0.27 (95% CI = 0.26, 0.28) in 2019 to *z* = 0.25 (95% CI = 0.23, 0.27) in 2016. Values for mango ranged from *z* = 0.43 (95% CI = 0.42, 0.45) in 2021 to *z* = 0.36 (95% CI = 0.35, 0.38) in 2016. Values for tamarind ranged from *z* = 0.1 (95% CI = 0.08, 0.12) in 2019 to *z* = 0.14 (95% CI = 0.08, 0.2) in 2015. Flowering onset typically occurred 4-6 weeks before mean flowering week and showed more clustering within species than flowering status (Fig 2). Mean flowering onset across the years ranged from week -3.27 in 2017 to 3.96 in 2019 in jackfruit, −0.7 in 2019 to 3.57 in 2016 in mango, −23.49 in 2018 to 25.37 in 2015 in tamarind (Fig 2b). Temporal clustering in flowering onset was less variable across the years; with tamarind having lower onset clustering than jackfruit and mango, during most years. Clustering within species ranged had a smaller range, lowest clustering in jackfruit was observed in 2017 with *z* = 0.49 (95% CI = 0.42, 0.07) and highest in 2019 with *z* = 0.51 (95% CI = 0.46, 0.38) (Fig 2b). For mango, lowest and highest clustering of onset was observed in 2019 with *z* = 0.53 (95% CI = 0.47, 0.1) and 2016 with *z* = 0.55 (95% CI = 0.44, 0.47) respectively (Fig 2b). For tamarind, lowest clustering was observed in 2018 with *z* = 0.24 (95% CI = 0.08, 0.12) and highest in 2015 with *z* = 0.44 (95% CI = 0.08, 0.2) (Fig 2b). There were no observable directional trends with time in timing or clustering of flowering.

**Figure 2:**
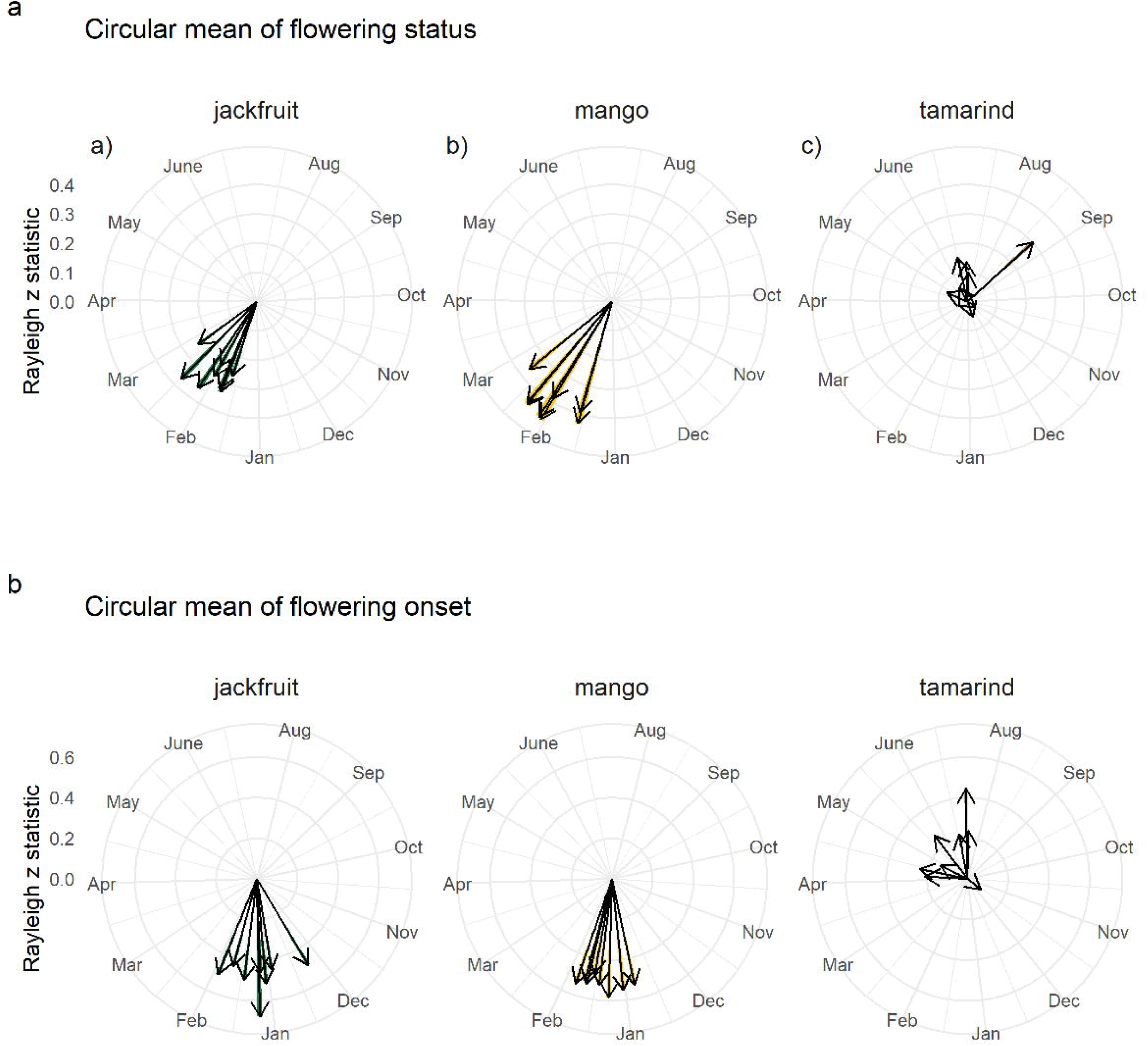
Mean date of flowering presence and flowering onset from 2014-2022 for the species of interest in Kerala. Bootstrapped estimates of flowering presence from weekly observations each year for a) jackfruit, b) mango, and c) tamarind. Bootstrapped estimates of flowering onset (first definitive observation of flowering on any tree) from weekly observations each year for d) jackfruit, e) mango, and f) tamarind. For a given year, arrow direction represents the mean week and arrow length represents z-score, a measure of degree of clustering from a bootstrapped Rayleigh test. See methods for details on how these were calculated.

Individual trees across all three species exhibited large variation in mean flowering onset week across the years as well as low consistency in the date of onset (Fig 3). Across all trees with observed true onset across the years, mean flowering onset week was distributed widely across the year for all three species (Fig 3a). Most trees of jackfruit showed a mean flowering onset week in the calendar year of -6, mango in week −12 and tamarind in week −20 (Fig 3a). However, trees showed large variation in onset week across years (Fig 3b). For trees with at least three years of onset observations, median standard deviation of flowering onset week was 6.91 for jackfruit, 6.39 for mango and 15.94 for tamarind.

**Figure 3:**
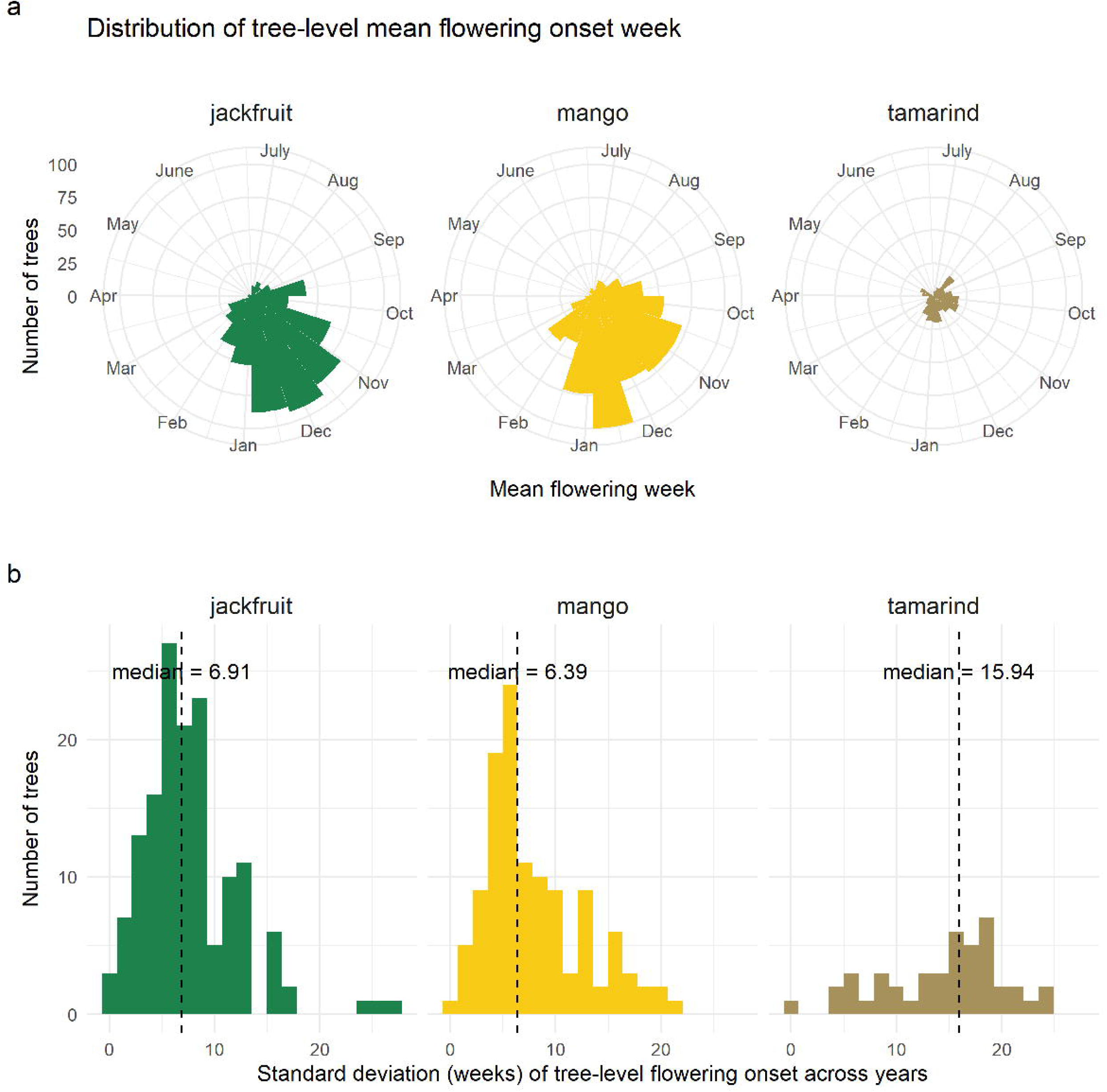
Variability in flowering onset across years for the same tree from 2014-2022. Distribution of mean week of flowering onset across years for trees with multiple onset observations for a) jackfruit, b) mango, and c) tamarind. Distribution of the standard deviation of flowering onset weeks for trees of a) jackfruit, b) mango, and c) tamarind.

### Proximate environmental cues for flowering onset on a tree

Flowering onset at the tree level across Kerala was predicted by minimum temperature, soil moisture and solar radiation but in varying magnitudes and directions across species (Fig 4). For jackfruit and mango, flowering onset was associated with lower minimum temperature and drier soils while for tamarind with higher minimum temperatures and wetter soils (Fig 4). Solar radiation had a weak positive effect on the probability of flowering in mango (Fig 4). GLMMs models of flowering onset had substantially better fits for jackfruit and mango than tamarind; the conditional R^2^ for the models, a metric of model fit including random effects, for jackfruit, mango and tamarind were 0.228, 0.242 and 0.131. Marginal R^2^, that only includes predictions using fixed effects, was 0.079, 0.102 and 0.026 for jackfruit, mango and tamarind, suggesting poor predictability from environmental variables alone, especially for tamarind. This suggests that fixed effects of environmental variation alone explained only a small proportion of the variation across flowering onset observations.

**Figure 4:**
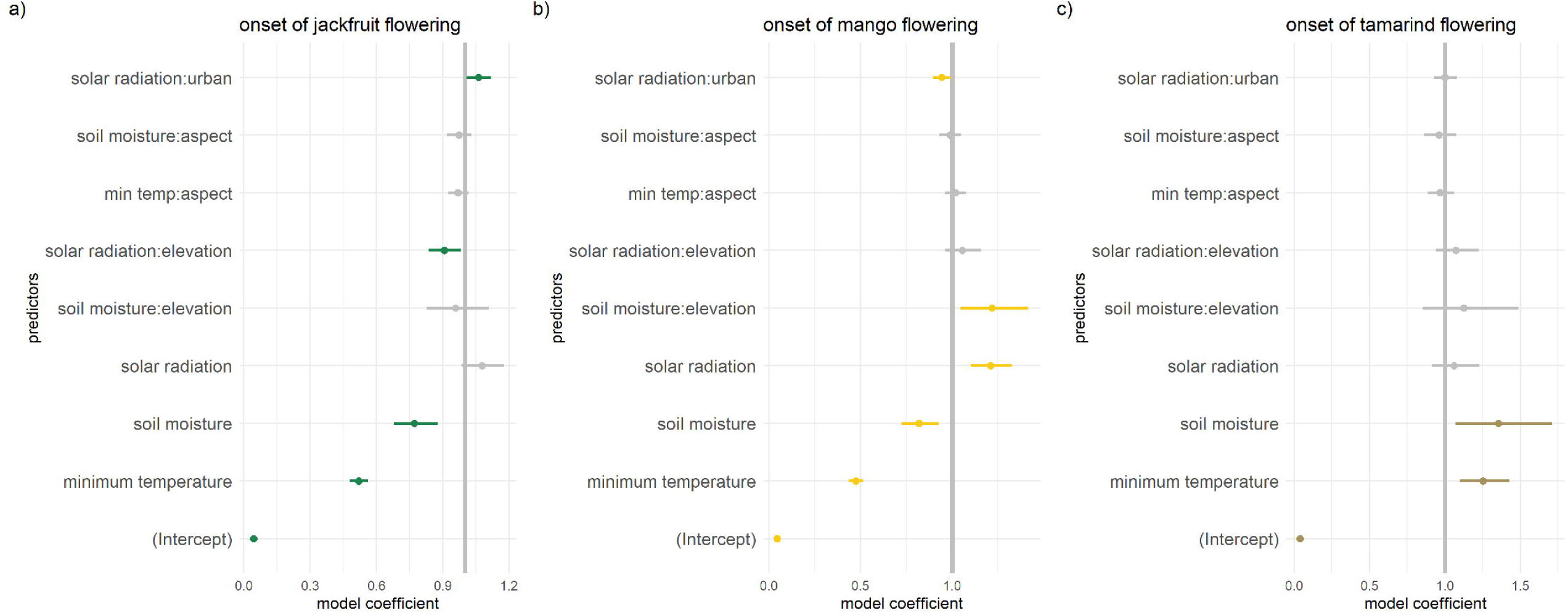
Model coefficients for flowering onset from GLMM models. Generalised linear mixed effect models (with a binomial family of errors) modelling flowering onset; i.e. the probability of observing flowering given no flowering at the previous timestep. Model includes most influential temperature variable, precipitation variable from the ML models and solar radiation. Model also includes all interactions barring known correlated combinations. Dots reporesent mean effect and the lines represent 95% confidence intervals of the estimate. The vertical grey line represents no-effect. Predictors with confidence intervals that do not overlap with the no-effect line are considered significant; others are represented with grey dots.

The ICC of the models were 0.162, 0.156 and 0.107 suggesting that random effects were not very clustered. To assess the predictive power of the model for out-of-bag data, we assessed the agreement between the predicted and observed values from the onset model for one year of the data that was withheld from modelling across 9 LOOCV runs. Coefficient across these runs were similar in magnitude and direction (Appendix S1: Section S6). Pearson’s correlation coefficients for predicted vs observed for jackfruit and mango across these runs 0.011, 0.031. The average mean square errors, respectively were 0.043, 0.041. F1 scores were 0.002, 0.007. The model was unable to predict flowering onset in tamarind with all predictions tending to non-flowering.

Elevation and degree of urbnaisation modified the influence of environmental factors on the onset of flowering in jackfruit and mango trees (Fig 4). Drier soils were associated with higher probability of flowering onset in mango, but this effect was stronger in lower elevations. In parallel, solar radiation increased the probability of flowering onset in mango trees, but this effect was dampened in more urbanised areas (Fig 4). In jackfruit trees, the influence of solar radiation as a cue for flowering onset was greater in lower elevation and more urban areas (Fig 4). However, main effects of solar radiation were not significant at the 90% confidence level, making interpretations of these interactions difficult. No interactions were significant for onset of flowering in tamarind trees. However, the model was a poor fit, and therefore, the possibility of significant interactions with more data cannot be ruled out.

### Spatiotemporal patterns in environmental variables

Locations across Kerala are experiencing heterogenous change in climatic conditions. Emerging hotspot analysis revealed spatiotemporal patterns in the three variables assessed, minimum temperature, soil moisture and solar radiation, in the time period when SeasonWatch data was collected (Fig 5). Aggregated across the state, these environmental variables did not show significant overall directional trend of change, defined as new or intensifying hotspot/coldspot, at the spatial scale assessed, from 2014-2022. Along the west to east elevational gradient in Kerala, minimum temperature had persistent hotspots in coastal locations, while inland sites towards the east were persistent cold spots, potentially due to higher elevation (Fig 5). Emerging hotspot analysis for soil moisture revealed a persistent hotspot in the south centre of the state. This hotspot corresponds to the eastern parts of the districts of Ernakulam and Kottayam, and the western parts of Idukki. For minimum temperature and soil moisture, pixels around the persistent hotspots, show varying and heterogenous trends in hotspots or coldspots. Mid-elevation sites in Kerala between the persistent coldspot and hotspot for minimum temperature have been classified as oscillating, sporadic or historical hot/coldspots, suggesting heterogenous change. Similarly for soil moisture, areas around the persistent hotspot show sporadic or oscillating hotspots. No directional patterns were discernible for solar radiation.

**Figure 5 -.**
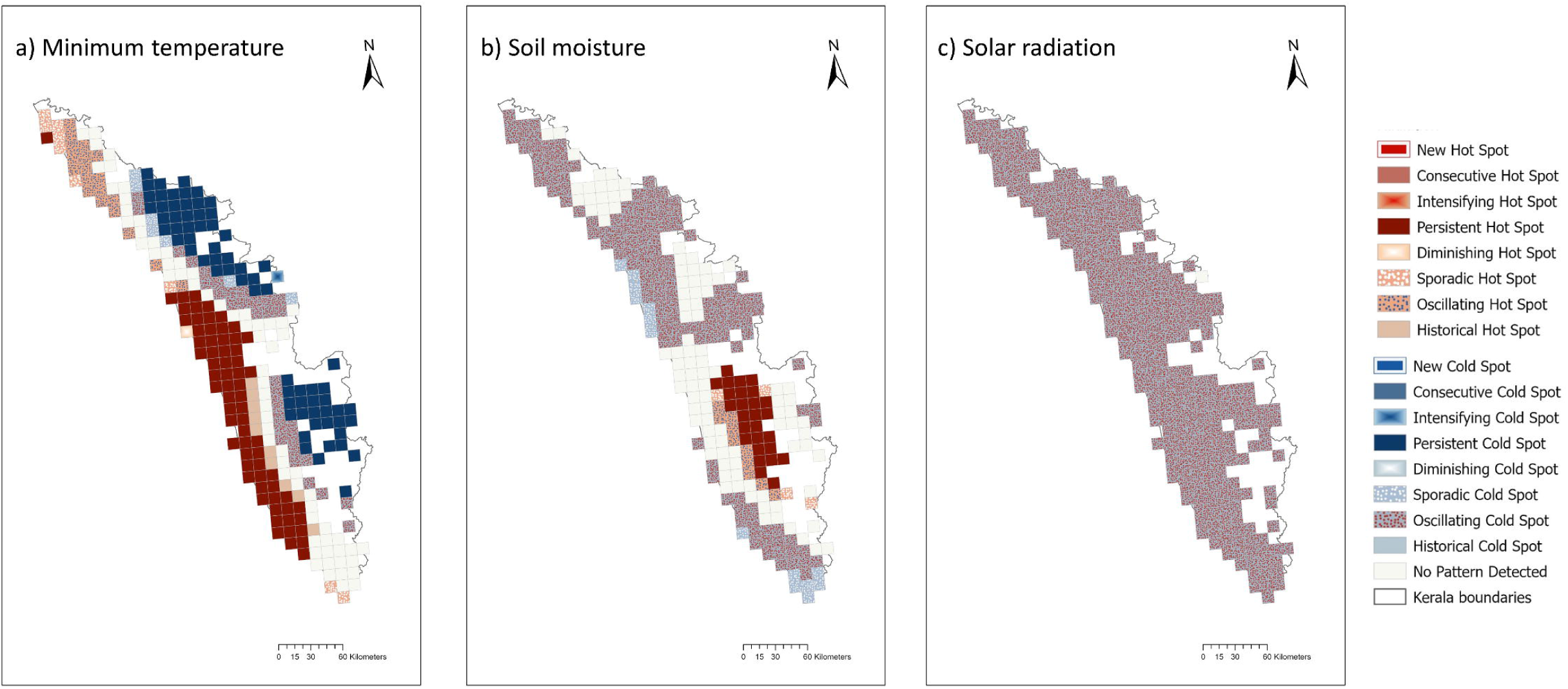
Hotspot analysis of environmental data using spatiotemporal data from 2014-2022. Hotspot analyses for a) minimum temperature within a fortnight, b) soil moisture within the fortnight and c) solar radiation within the fortnight within each grid of 11 km x 11 km spatial resolution

## Discussion

Altered temperature and precipitation patterns are expected to affect phenological responses of tropical tree species, but their variability across large spatial scales is rarely quantified and poorly understood. We used a unique citizen science dataset from the southwestern state of Kerala in India (Fig 1), unparalleled in scale for the tropics, to examine seasonality in flowering in common tropical tree species and the extent and correlates of interannual and interindividual variability. Our results show that over 9 years of weekly observations, timing of flowering status and flowering onset across three species shows 4-6 weeks of variation across years at the population level (Fig 2). This interannual variation in the timing of flowering was not mirrored in temporal clustering flowering observed at the population level, which remained relatively constant, but was reflected in individual trees that showed substantial interannual variation in timing of flowering onset(Fig 3). We show that some of this individual-level variation in phenology of flowering onset across the years is associated with recent, local climatic conditions, such as soil moisture and minimum temperature, the effects of which can be different along elevation and urbanisation gradients (Fig 4). These environmental cues associated with phenology of these common trees, however, are changing across Kerala (Fig 5), with potential consequences for phenology and associated ecosystem services. Our results show the potential of citizen science datasets to provide crucial insights at spatial and temporal scales relevant for adaptive ecosystem management.

Our use of citizen science data and associated temporal data gaps limits the use of some common methods like Fourier transformation to disentangle seasonal patterns from long-term drivers at the individual tree level. Cohort-based long-term monitoring of tree populations has led to important understanding of the influence of seasonal precipitation and interannual variability through ENSO events, droughts etc. on reproductive phenology of trees in tropical sites (Wright et al. 1999, Harrison 2000, Sakai and Kitajima 2019). Given the lack in uninterrupted timeseries at the individual tree level, we were unable to disentangle individual variation in phenology from temporal patterns for long timescale. However, our study spans nine years of observations, more than the minimum recommended 6 years for tropical species (Bush et al. 2018), and the spatial extent of the dataset is much larger compared to the limitations of long-term cohort monitoring (often restricted to a site) allowing us to extend inferences to large spatial scales. Examining flowering observations across a wide variation in habitats also revealed that trees in high elevation or urban areas respond differently to environmental cues, adding to prevalent understanding of the contextual nature of flowering cues (Neil and Wu 2006, Babweteera et al. 2018, Kabano et al. 2020). Such accumulating broad-scale datasets led by citizen scientists could be important complements to finer scale long-term expert-led monitoring to understand shifting phenological baselines across species ranges.

Extending our results across tropical trees and landscapes poses certain limitations because of some broad ecological and functional similarities in the three species studied. All three species studied were dioecious, evergreen, widespread, insect-pollinated, bear fleshy fruits, and cultivated in home gardens (Table S1). Notably, jackfruit trees in the dataset showed high tendencies for multiple instances of flowering onset within a year, and tamarind showed low seasonality in flowering overall (Fig 2). Mixed models of flowering onset showed that environmental predictors explained only ∼10% of variation in probability of switch to flowering status, while including random effects of sites explained ∼20% of the variation across all three species. Since these are cultivated species, the differences in varieties among individuals has a known influence on phenological variation (Makhmale et al. 2016, Singh et al. 2021).

Applicability of these inferences across diverse tropical species with different leaf and reproductive phenology could pose challenges. Reproductive strategies in tropical trees are often diverse and may have long-term evolutionary function in addition to short-term ecological response (Singh and Kushwaha 2006). Borchert et al. (2004) showed that flowering seasonality in tropical species can be plastic across biome ranges, with greater periodicity in the drier forests across species. We show that even in low latitudes with low seasonality, flowering status and flowering onset within species show strong periodicity and interannual variation associated with climatic variables. Given the phylogenetic distinctiveness among the three species, we are encouraged to conclude that these conclusions could apply over a broader set of species.

Our results become pertinent under existing and future climatic variability in the tropics, and the potential impacts on the reproductive phenology of commercially important tropical trees.

Beyond the timescales of our study, much of the global tropics, is projected to move into a temperature and precipitation regime that has no analog globally, making it difficult to predict and plan for ecosystem responses (Dahinden et al. 2017). Across the Indian subcontinent, precipitation and temperature variability has increased in the past century and is projected to continue to increase (Power et al. 2013, Dahinden et al. 2017, Zhang and Fueglistaler 2019). Moreover, altered climatic processes such as the El Nino Southern Oscillation and Indian Ocean dipole are affecting weather with consequences for Indian ecosystems (Revadekar et al. 2009, Wang and Wang 2014, Gadgil and Francis 2016). However, climate change predictions and action for the tropics are still largely at coarse spatial scales, making local forecasting and mitigation difficult (Dessai et al. 2009, Shepherd and Sobel 2020). Across Kerala, our results from emerging hotspot analyses show spatiotemporal patterning in climate variables associated with flowering phenology in common tree species (Fig 5). An important future direction for phenological studies would be to understand intraspecific variation across space and forecast phenological responses in these novel climates.

Our study using repeat citizen-science data and individual based modelling at spatial and temporal scales of climate modelling, represents a significant first step in combining tropical ecology with climate science beyond broad generalisations. Despite recent remote sensing advances that aid large-scale and repeat measurement of ecosystem structure and change, measurement of tropical reproductive phenology remains challenging (Davis et al. 2022, Lee et al. 2023). Species may have inconspicuous flowers and fruits or these phenophases may coexist with leaves, requiring repeat on-ground measurements that are extremely difficult to organise at the large-scale. Citizen science may be one of the few ways to acquire such data, and we have leveraged such a unique dataset compiled by SeasonWatch to predict the onset of a reproductive phenophase (Abernethy et al. 2018, McDonough MacKenzie et al. 2020, Ramaswami et al. 2021). Broadly, we demonstrate the potential of citizen science observations in improving the state of our knowledge in climate science and infusing local ecology into climate-adaptive management. Predictive modelling informed by variation among individuals and sites is a crucial next step to phenology research as spatiotemporal patterns in climate variables change across tropical landscapes. Our study thus demonstrates the potential of citizen science in filling data gaps from expert-led site-specific monitoring and providing predictions at the large-scale to aid locally relevant climate policies.

## Acknowledgements

The authors would like to acknowledge the SeasonWatch team for support during this work, especially Mohammed Nizar, who helped in the manual validation of parts of the data. We also thank Karthik T, Anand M O, Suhel Quader and the NCF EPE team for discussions on data management, analyses and interpretations. SW is grateful for the support of the Wipro Foundation for their continued efforts.

## Research ethics

The study did not deal with human subjects or data with personal identifiers. For SeasonWatch data, we followed ethical protocol that was agreed on by the SeasonWatch Citizen Scientist Network.

## Author contributions

Funding for this work was acquired by GR and NA, while the study was conceptualised by GR and KA. Phenology data was collected by the SeasonWatch Citizen Scientist Network. KA cleaned and validated the phenology data while JM accessed and collated the geospatial data. KA, JM, CL and RR performed the analysis led by KA and supervised by GR and NA, with administrative support from HT. KA visualised the data and wrote the first draft; KA, JM and GR edited the manuscript with support from all authors.

