## Supplementary Appendix for "Long-term citizen science data reveals environmental correlates of tropical tree flowering at the regional scale"

***Journal:*** Ecological Applications

***Authors:*** Krishna Anujan[1,2]*, Jacob Mardian[3], Carina Luo[3], Ramraj Rajakumar[3], Hana Tasic[3], Nadia Akseer[3], Geetha Ramaswami[1]

***Affiliations:***
[1] SeasonWatch, Nature Conservation Foundation, 1311, "Amritha", 12th Main, Vijayanagar 1st Stage, Mysore 570017, India
[2] Department of Ecology, Evolution and Environmental Biology, Columbia University in the City of New York, 1110 Amsterdam Ave, New York, NY 10025, United States
[3] Modern Scientist Global, 110 James Street, Suite 101, St. Catharines, Ontario L2R 7E8, Canada

#### Section S1. Species descriptions and SeasonWatch observations

Table S1. Species descriptions.**Taxonomic description, appearance of tree and reproductive parts, phenology in Kerala and description of the SeasonWatch dataset for jackfruit, mango and tamarind

| | Jackfruit | Mango | Tamarind |
|---|---|---|---|
| Accepted Scientific name | Artocarpus heterophyllus | Mangifera indica | Tamarindus indica |
| Name in Malayalam | Pilavu/Plavu (tree), chakka (fruit) | Maavu/Moochi (tree), manga (fruit) | Puli, Valan puli, Kolpuli |
| Family | Moraceae | Anacardiaceae | Fabaceae |
| Description (Trees of Delhi) | A handsome evergreen tree with glossy leaves up to 20 m tall in ideal conditions. Cultivated for its misshapen fruits - largest edible fruit in the world | Semi-evergreen trees with long, glossy leaves that can grow up to 35 m in favourable sites. Bark is grey-brown and rough with shallow cracks. | Large, handsome, long-lived tree with a squat trunk and a shady crown. Introduced to India from E Africa, but virtually indigenous. The bark is dark grey, rough with shallow fissures |
| Flower description | Male and female separate, on the same tree. Male flowers tiny, in clusters emerging from new branchlets; female flowers densely crowded on old branches or directly from the trunk | Mango flowers are tiny, yellowish-green, strongly scented, in huge, branched clusters. Most flowers are male and the rest bisexual. | Small, with 3 unequal-sized yellow petals veined with red; flower cup has 4 creamy segments |
| Fruit description | Gigantic, lumpy, barrel- or pear-shaped; pollinated by small flies and beetles | Fruit is smooth-skinned, waxy and in various colours depending on the variety. Flesh is fibrous or pulpy and the stone flattened, kidney-shaped | Beanlike pod bulging over the seeds, up to 20 cm long; green at first, ripening cinnamon-brown. Young pod is covered with a downy felt. |
| Common varieties in Kerala | Varikka, koozha, etc | Muvandan, chandrakkaran, etc | |
| Uses | Timber, fruit | Fruit | Fruit, timber |
| Fruiting and flowering months in Kerala | November to April | January to May | September to April |

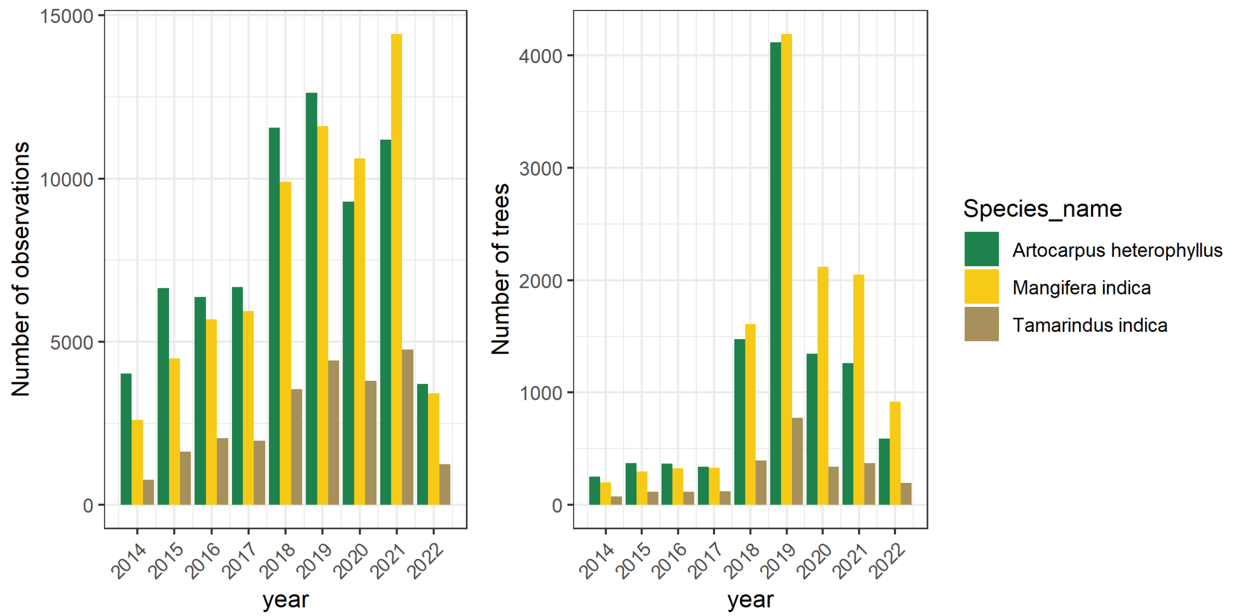

Figure S1. Data distribution of all SeasonWatch observations for the top three species in Kerala across the years. a) Total number of observations including regular and casual b) Number of trees in these observations

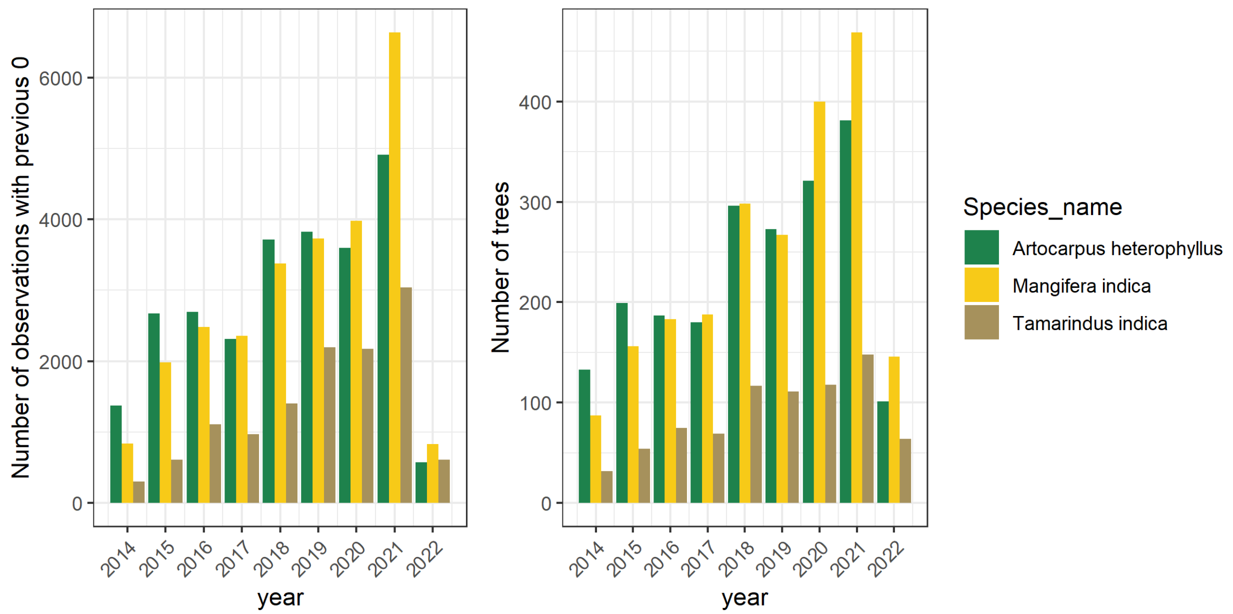

Figure S2. Data distribution of SeasonWatch observations with a 0 flowering observation in the previous week for the top three species in Kerala across the years. a) Total number of observations (includes only regular because casual observations do not have previous week's observation) b) Number of trees in these observations

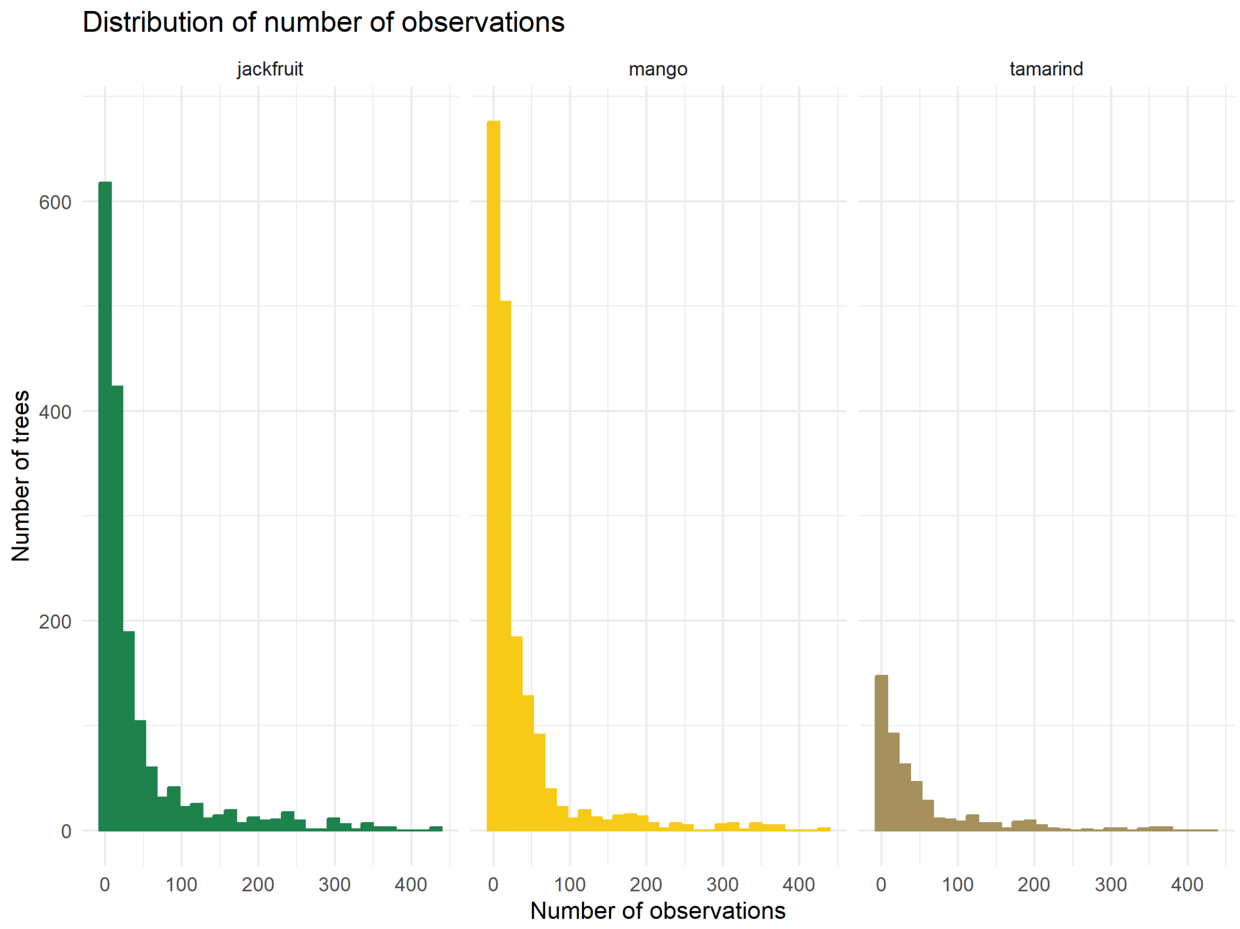

Figure S3. Distribution of regular observations of the top three species in Kerala across the years.

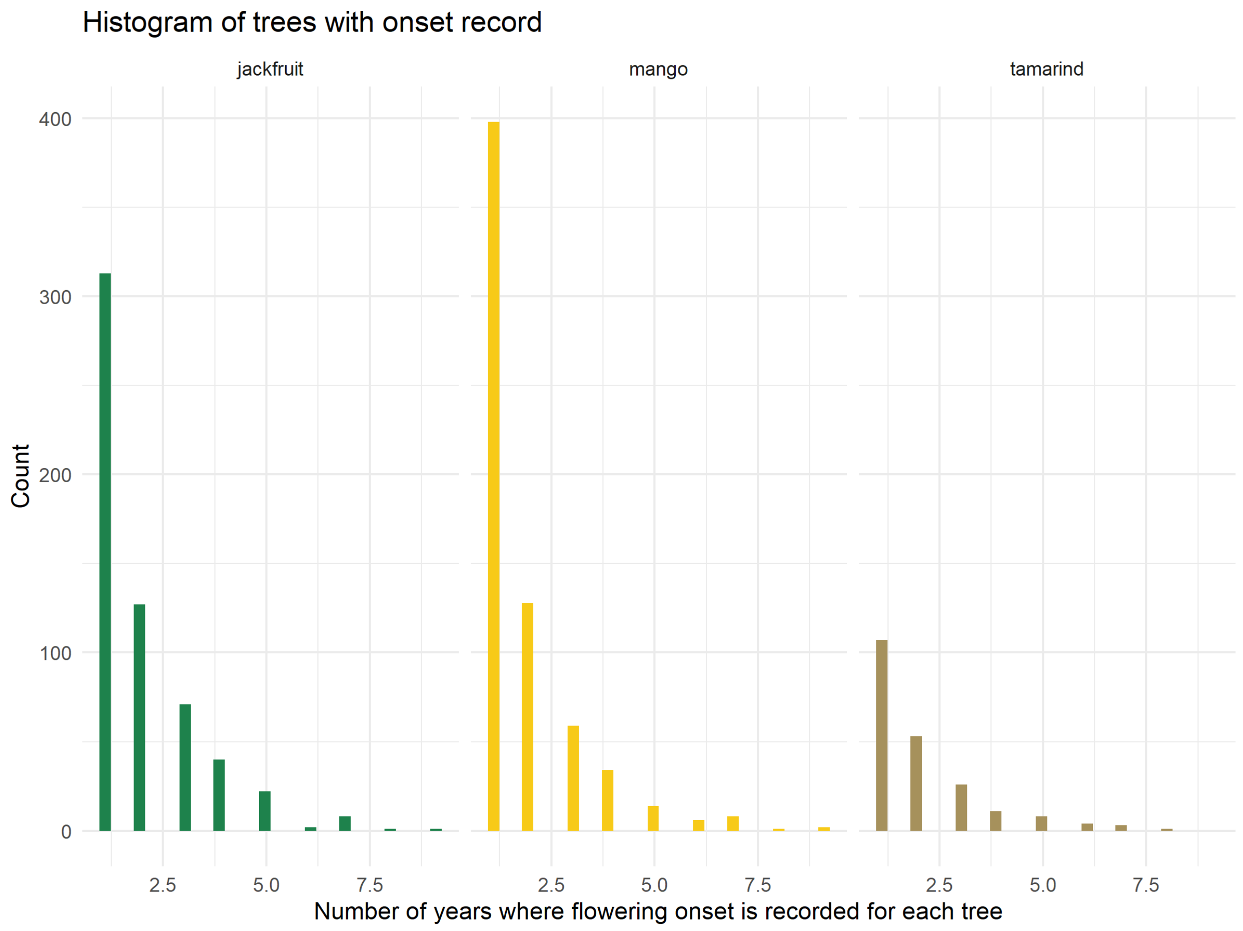

Figure S4. Number of years with observed onset record for each tree

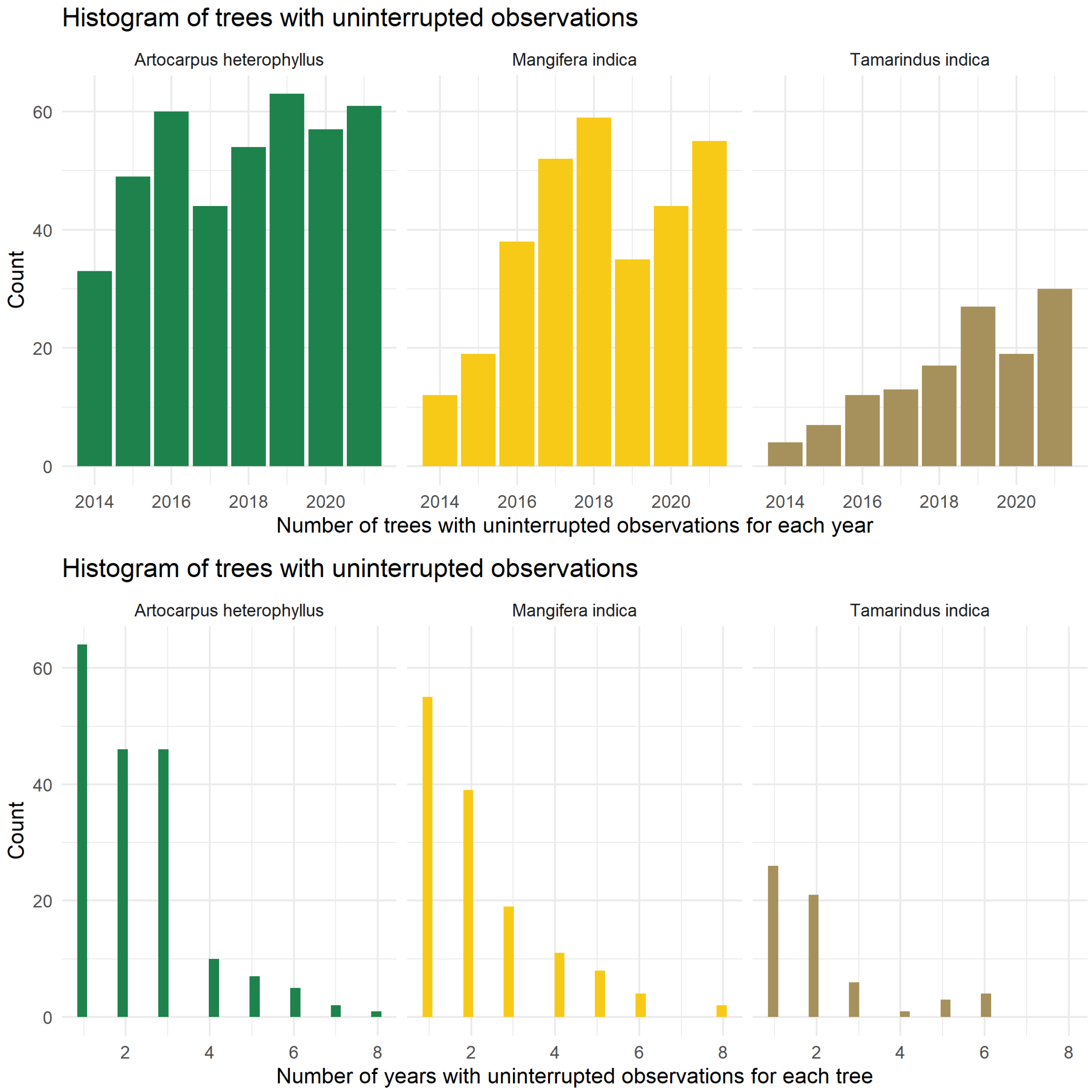

Figure S5. Distribution of trees with uninterrupted flowering record

#### Section S2. Supplementary methods for data cleaning

##### Geographic location of points

To validate the geographic location of the data points, we included all the points that were already in Kerala as per the geographic location. From the remaining points, for any that had no location information, we excluded those that were single observations of trees, because of no alternate way to fill these. For regular observations with unconfirmed locations, we filled them with lat/longs from the same tree if available in later observations, or the average lat/long from other trees reported by the same user. As majority of the users were school students observing trees on foot, all regular trees observed by the same user tend to be highly clustered ($< 1$ km), allowing us to reliably fill an average lat/long from other trees when no observation of a tree had a lat/long. Finally, if no trees by the user had accurate lat/long values and if they were by users who can be located through Google maps (schools or other institutions), we used a geocoding software on Google Sheets, Awesome Table Geocode, to fill these values. We then manually validated these with help from the field coordinator for Kerala. We excluded observations that could not be filled or verified by these methods.

#### Section S3. Environmental variable methods

##### Climate Data Column Descriptions

All climate data was calculated on the daily time step for 0.1° pixels and aggregated to the fortnightly, monthly, and quarterly timesteps. A summary of the daily climate data is shown below:
* PCP = Total daily precipitation (in meters)
* MinTemp = Minimum daily temperature (in Kelvin)
* MeanTemp = Average daily temperature (in Kelvin)
* MaxTemp = Maximum daily temperature (in Kelvin)
* MeanSM = Average daily soil moisture (the top 1m of the soil column in volumetric soil water content: m3/m3)
* SolarRad = Total surface net solar radiation (in J/m2)
* DryDays = Maximum consecutive days with PCP < 1mm (trace amounts are a common artefact in climate reanalysis data)

For more information about these variables (except dry days, which was calculated from PCP), see https://developers.google.com/earth-engine/datasets/catalog/ECMWF_ERA5_LAND_HOURLY

Each variable has a fortnightly, monthly, and quarterly aggregation. Precipitation and Solar Radiation are accumulated totals (i.e., sum) while the temperature and soil moisture variables are averages. Columns abbreviated "FN" indicate fortnightly, "MN" indicates monthly and "QR" indicates quarterly. For example, "FNMean_MaxTemp" indicates that the mean Maximum Daily Temperature value was taken for the previous fortnight. "QrSum_PCP" indicates that the sum of daily total precipitation was taken for the previous quarter.

Static Data Column Descriptions * light_vals = Static variable. NASA satellite-derived lights dataset as a proxy for urban density
* elev = Static variable. Elevation above mean sea level derived from 30m SRTM
* slope = Static variable. Rise/run of elevation dataset
* aspect = Static variable. Direction of slope

Observation-Level Data

The dataset includes the Grid_ID and Fortnight they belong to, the observation ID, date of observation, species name, latitude, longitude, the response variables (fl_binary, fl_intensity) and the climate indicators. FN: Previous 14 days before date of observation MN: Previous 30 days before date of observation Qr: Previous 90 days before date of observation

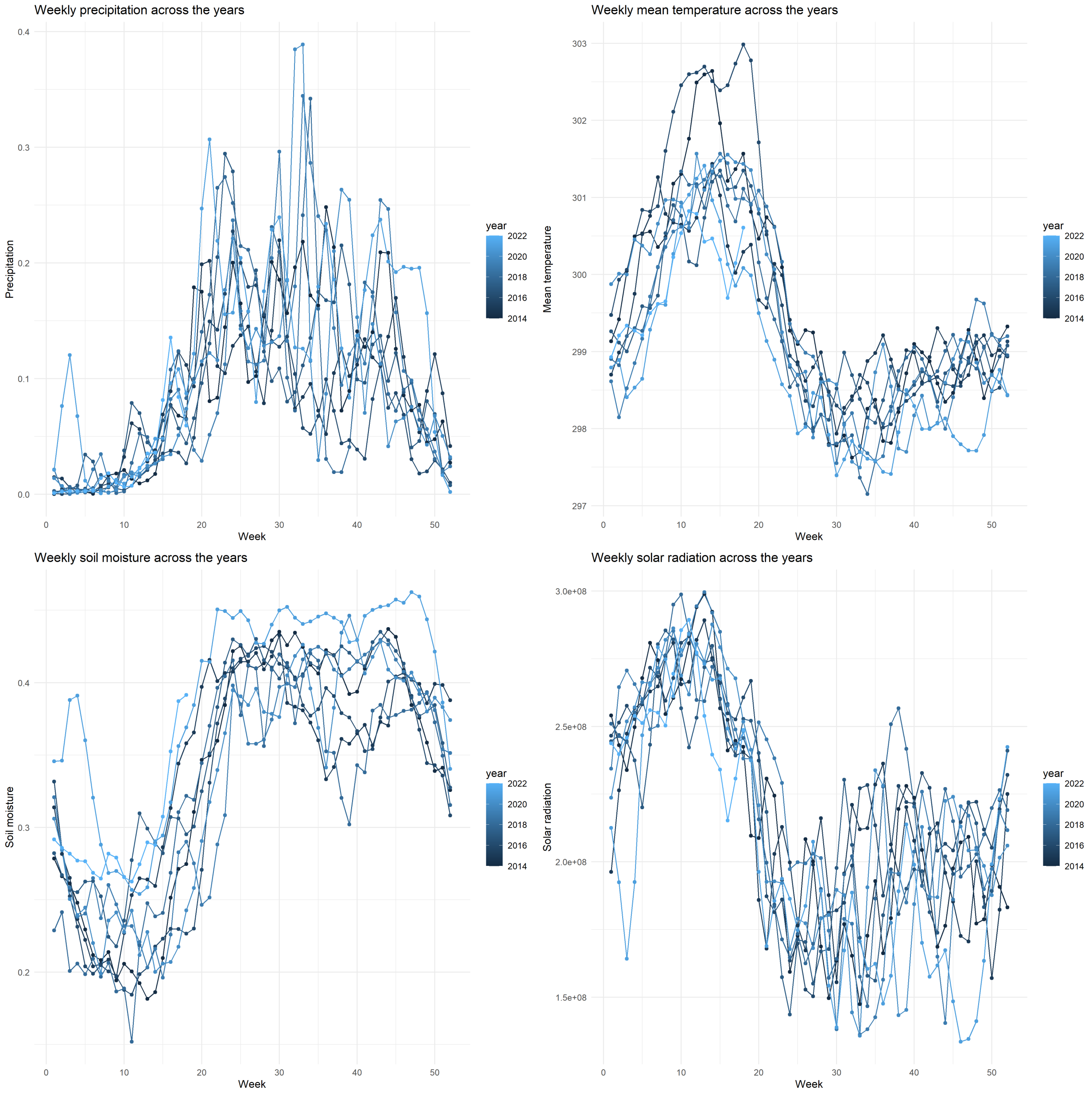

Figure S6. Mean weekly climate observations across locations where trees have been observed in Kerala through SeasonWatch

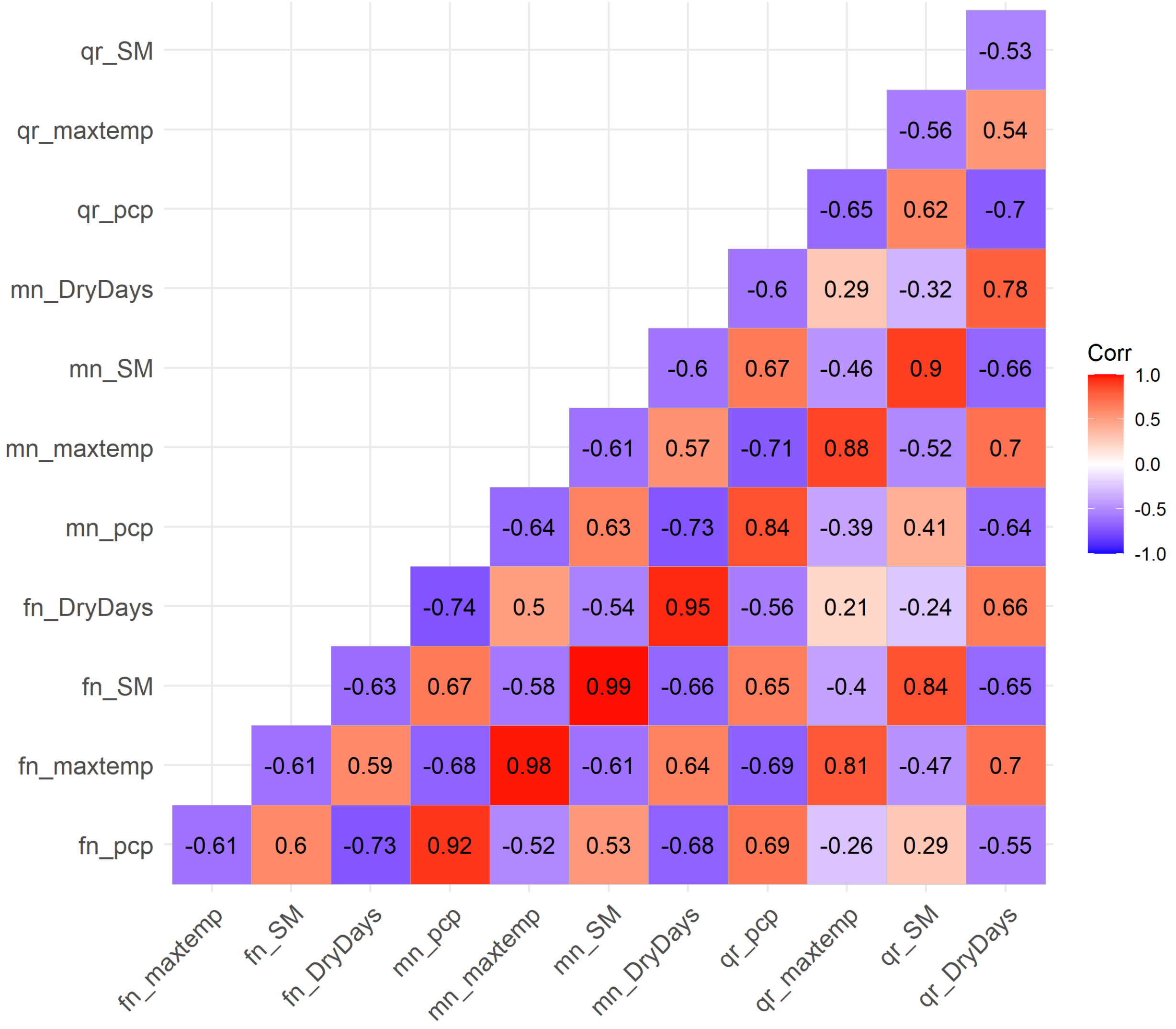

Figure S7. Correlations between environmental variables at different timescales

#### Section S4. Supplementary methods and results for ML modelling

For the entire set of observations, we used Random Forest models to classify the flowering occurrence observations based on environmental predictors. Random Forest classifier is a supervised learning algorithm that uses an ensemble of decision trees using majority voting to classify datasets into two categories; here the presence/absence of flowering (Breiman 2001). Random Forest classifiers have been used widely in ecological data analysis because of their flexibility and accuracy (Cutler et al. 2007). This approach allows for predicting the probability of observing flowering in a tree given recent environmental conditions.

For each of the three species, we ran separate models to classify flowering status according to the following model:

$$phenophase_{l,t} \sim precipitation_{l,t} + maximumtemperature_{l,t} + minimumtemperature_{l,t} + meantemperature_{l,t} + drydays_{l,t} + so$$

Where precipitation, temperature, dry days, soil moisture, solar radiation are temporal predictors for the fortnight before the observation at location $l$ and time $t$. We ran these models in R using the package *randomForest* (Liaw and Wiener 2002) for 500 decision trees. Tree-based methods are fairly insensitive to collinearity among predictors, allowing us to compare the full-range of temperature and precipitation variables simultaneously, as well as the full dataset, including repeat observations for the same tree.

We calculated variable importance for each model using the Mean Decrease in Accuracy (MDA) metric which removes a predictor from the model, re-trains the model to make predictions, and evaluates the decrease in accuracy.

Machine learning models with spatiotemporal predictors were able to classify flowering phenophases for the three species with substantial precision. The accuracy of a binary model by random chance would be 50%, leading to an F1 score of 0.5; models with higher F1 scores than randomly expected have predictive value. With 500 runs each, Random Forest models were able to predict the presence/absence of flowering in jackfruit, mango, and tamarind trees given the environmental conditions of the previous fortnight with F1 scores of 0.59, 0.62, and 0.26, respectively. These performance metrics describe the ability of the models to predict tree phenophases for an unobserved year based on environmental variables alone. We note that the distribution between flowering and no-flowering observations was about 2:1, potentially influencing the F1 score.

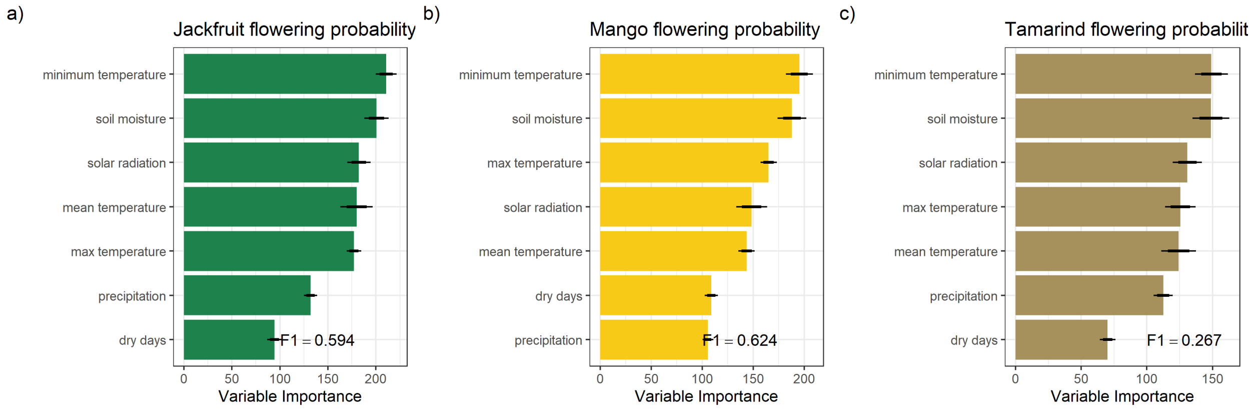

Random forest models identified environmental variables strongly correlated with flowering for each of the species. The mean importance of each of the factors, calculated using MDA, was different across species. For all three species, minimum temperature in the preceding fortnight was the most important predictor of flowering status. Among precipitation variables, soil moisture in the preceding fortnight had the strongest influence in classifying flowering status. Solar radiation also ranked among the top three variables for all three species. Across the 9 LOOCV runs, the relative importance of the variables had some variation, but total precipitation and the number of consecutive dry in the previous fortnight consistently were ranked lowest across runs in all species. Some of the variation in importance was also potentially influenced by the difference in number of observations across the years, with 2014 being the lowest sample.

Table S2. Variable importance across each of the LOOCV models for jackfruit

| | mean temperature | minimum temperature | max temperature | solar radiation | precipitation | dry days | soil moisture |
|---|---|---|---|---|---|---|---|
| 2014 | 194.2509 | 214.5788 | 179.2915 | 186.6246 | 135.1327 | 95.93955 | 194.1267 |
| 2015 | 179.5465 | 209.1443 | 188.5371 | 163.9275 | 126.3947 | 94.94915 | 198.4492 |
| 2016 | 190.2715 | 219.5325 | 175.8974 | 186.1020 | 132.2170 | 89.85891 | 202.5824 |
| 2017 | 165.7244 | 216.9539 | 172.4679 | 188.7910 | 135.5556 | 97.61727 | 215.0707 |
| 2018 | 163.4843 | 214.7208 | 175.2011 | 183.9784 | 127.4882 | 102.55036 | 195.5013 |
| 2019 | 184.4766 | 208.3534 | 176.8716 | 183.7102 | 139.4553 | 89.46958 | 207.0071 |
| 2020 | 184.7028 | 204.5251 | 174.5972 | 179.6201 | 133.1122 | 96.71695 | 207.2555 |
| 2021 | 173.7381 | 197.3297 | 173.6899 | 180.2438 | 130.7285 | 87.91403 | 189.8676 |
| 2022 | 182.7083 | 210.4814 | 177.0990 | 187.7085 | 127.7086 | 92.99067 | 195.6885 |

Table S3. Variable importance across each of the LOOCV models for mango

| | mean temperature | minimum temperature | max temperature | solar radiation | precipitation | dry days | soil moisture |
|---|---|---|---|---|---|---|---|
| 2014 | 134.4052 | 197.1949 | 164.8218 | 156.2773 | 104.72479 | 110.60015 | 202.8316 |
| 2015 | 144.9679 | 198.7177 | 163.3524 | 141.5846 | 109.64898 | 114.76737 | 180.4693 |
| 2016 | 144.6262 | 189.7379 | 155.3212 | 148.4978 | 107.66133 | 108.66425 | 200.1551 |
| 2017 | 144.4997 | 203.6345 | 167.7915 | 163.6967 | 110.57972 | 111.81441 | 187.1588 |
| 2018 | 138.2928 | 186.8528 | 163.5042 | 144.1055 | 103.84531 | 110.47167 | 178.7136 |
| 2019 | 140.8394 | 193.5883 | 163.4009 | 143.2801 | 99.64859 | 108.26231 | 186.1939 |
| 2020 | 144.5293 | 188.2500 | 169.6981 | 138.2137 | 106.36116 | 107.26544 | 191.2922 |
| 2021 | 148.4355 | 187.6162 | 163.8984 | 139.5365 | 104.15356 | 99.78337 | 178.4911 |
| 2022 | 150.8697 | 211.2650 | 173.2673 | 160.6331 | 103.66976 | 108.27752 | 185.9629 |

Table S4. Variable importance across each of the LOOCV models for tamarind

| | mean temperature | minimum temperature | max temperature | solar radiation | precipitation | dry days | soil moisture |
|---|---|---|---|---|---|---|---|
| 2014 | 125.3954 | 157.0129 | 123.5367 | 137.5659 | 112.1205 | 73.43858 | 161.9501 |
| 2015 | 112.7048 | 134.8631 | 119.5150 | 127.1199 | 108.4221 | 71.68865 | 145.6994 |
| 2016 | 133.7776 | 161.3472 | 123.6452 | 131.0377 | 109.8208 | 69.80641 | 152.8163 |
| 2017 | 129.6894 | 149.9381 | 122.4582 | 129.9581 | 112.6012 | 71.56924 | 154.0277 |
| 2018 | 112.3716 | 150.3709 | 129.8349 | 117.4087 | 105.5565 | 69.48091 | 149.1446 |
| 2019 | 117.0161 | 152.5773 | 114.9969 | 126.8927 | 117.6458 | 67.53726 | 144.7113 |
| 2020 | 124.7591 | 143.0515 | 131.3704 | 135.2399 | 119.6935 | 75.25158 | 131.8492 |
| 2021 | 131.1785 | 148.6381 | 124.0858 | 131.2129 | 110.6177 | 62.53390 | 155.7232 |
| 2022 | 131.0931 | 144.1736 | 140.0442 | 141.4173 | 116.0824 | 69.89865 | 143.3540 |

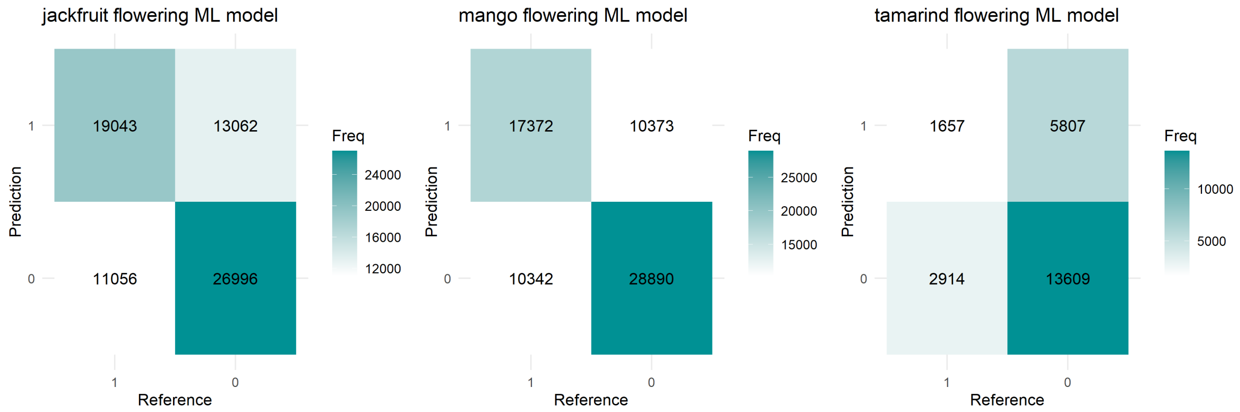

Figure S8. Confusion matrices with Random Forest models

#### Section S5. Supplementary results for GLMMs

Table S5. Model coefficients for flowering onset in jackfruit

### Fixed Effects

| Parameter | Log-Odds | SE | 95% CI | z | p |
|---|---|---|---|---|---|
| (Intercept) | -3.13 | 0.11 | (-3.35, -2.92) | -28.58 | < .001 |
| min temp | -0.65 | 0.05 | (-0.75, -0.56) | -13.62 | < .001 |
| soil moisture | -0.26 | 0.08 | (-0.41, -0.11) | -3.30 | < .001 |
| solar rad | 0.07 | 0.05 | (-0.03, 0.18) | 1.37 | 0.169 |
| elev cs | -0.17 | 0.06 | (-0.29, -0.05) | -2.73 | 0.006 |
| aspect cs | -0.08 | 0.04 | (-0.16, 6.07e-03) | -1.82 | 0.069 |
| lights cs | 0.06 | 0.06 | (-0.06, 0.18) | 0.92 | 0.356 |
| soil moisture × elev cs | -0.04 | 0.09 | (-0.22, 0.13) | -0.49 | 0.622 |
| solar rad × elev cs | -0.10 | 0.05 | (-0.19, -3.39e-03) | -2.03 | 0.042 |
| min temp × aspect cs | -0.03 | 0.03 | (-0.09, 0.03) | -1.07 | 0.285 |
| soil moisture × aspect cs | -0.03 | 0.04 | (-0.10, 0.04) | -0.78 | 0.433 |
| solar rad × lights cs | 0.06 | 0.03 | (-1.77e-03, 0.12) | 1.90 | 0.057 |

### Random Effects

| Parameter | Coefficient |
|---|---|
| SD (Intercept: Grid_ID) | 0.80 |

Table S6. Model coefficients for flowering onset in mango

### Fixed Effects

| Parameter | Log-Odds | SE | 95% CI | z | p |
|---|---|---|---|---|---|
| (Intercept) | -3.08 | 0.10 | (-3.28, -2.88) | -30.62 | < .001 |
| min temp | -0.75 | 0.05 | (-0.85, -0.65) | -14.59 | < .001 |
| soil moisture | -0.20 | 0.08 | (-0.35, -0.05) | -2.65 | 0.008 |
| solar rad | 0.19 | 0.06 | (0.08, 0.30) | 3.40 | < .001 |
| elev cs | -0.34 | 0.07 | (-0.48, -0.19) | -4.60 | < .001 |
| aspect cs | -0.03 | 0.04 | (-0.12, 0.06) | -0.71 | 0.478 |
| lights cs | 0.16 | 0.06 | (0.05, 0.27) | 2.87 | 0.004 |
| soil moisture × elev cs | 0.20 | 0.09 | (0.02, 0.38) | 2.14 | 0.032 |
| solar rad × elev cs | 0.05 | 0.06 | (-0.06, 0.16) | 0.97 | 0.334 |
| min temp × aspect cs | 0.02 | 0.03 | (-0.05, 0.08) | 0.51 | 0.609 |
| soil moisture × aspect cs | -0.01 | 0.04 | (-0.08, 0.06) | -0.31 | 0.754 |
| solar rad × lights cs | -0.06 | 0.03 | (-0.12, 2.38e-03) | -1.88 | 0.060 |

### Random Effects

| Parameter | Coefficient |
|---|---|
| SD (Intercept: Grid_ID) | 0.78 |

Table S7. Model coefficients for flowering onset in tamarind

### Fixed Effects

| Parameter | Log-Odds | SE | 95% CI | z | p |
|---|---|---|---|---|---|
| (Intercept) | -3.28 | 0.13 | (-3.53, -3.02) | -25.21 | < .001 |
| min temp | 0.23 | 0.08 | (0.07, 0.38) | 2.82 | 0.005 |
| soil moisture | 0.30 | 0.14 | (0.03, 0.58) | 2.14 | 0.032 |
| solar rad | 0.06 | 0.09 | (-0.12, 0.24) | 0.65 | 0.517 |
| elev cs | -0.05 | 0.16 | (-0.36, 0.26) | -0.32 | 0.752 |
| aspect cs | -0.08 | 0.07 | (-0.22, 0.07) | -1.02 | 0.306 |
| lights cs | -8.52e-04 | 0.07 | (-0.15, 0.14) | -0.01 | 0.991 |
| soil moisture × elev cs | 0.12 | 0.17 | (-0.21, 0.45) | 0.71 | 0.479 |
| solar rad × elev cs | 0.07 | 0.08 | (-0.09, 0.23) | 0.89 | 0.373 |
| min temp × aspect cs | -0.03 | 0.06 | (-0.14, 0.08) | -0.58 | 0.562 |
| soil moisture × aspect cs | -0.04 | 0.07 | (-0.17, 0.09) | -0.58 | 0.563 |
| solar rad × lights cs | 2.35e-04 | 0.05 | (-0.09, 0.09) | 4.96e-03 | 0.996 |

### Random Effects

| Parameter | Coefficient |
|---|---|
| SD (Intercept: Grid_ID) | 0.63 |

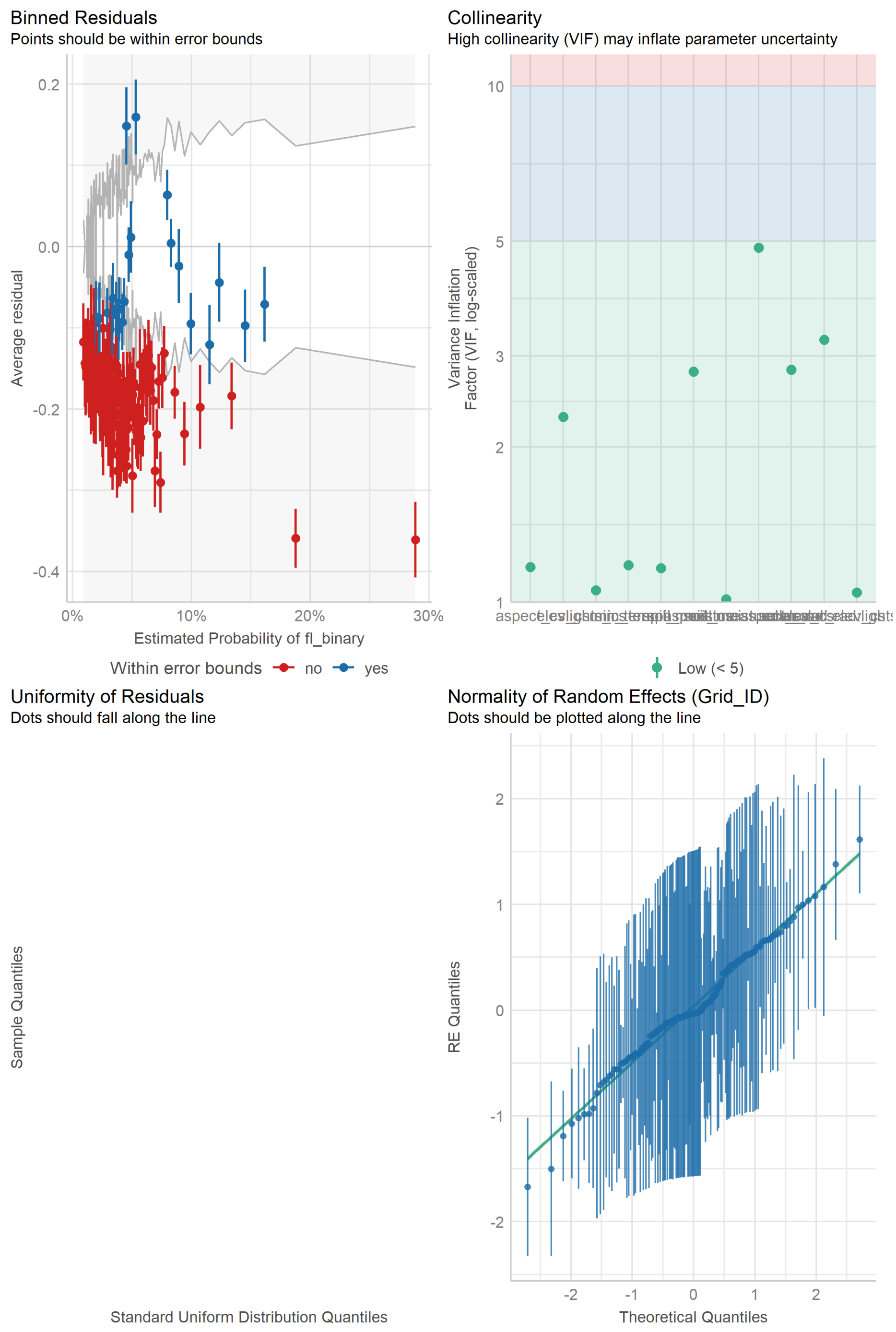

Figure S9. GLMM diagnostics for onset of flowering in jackfruit

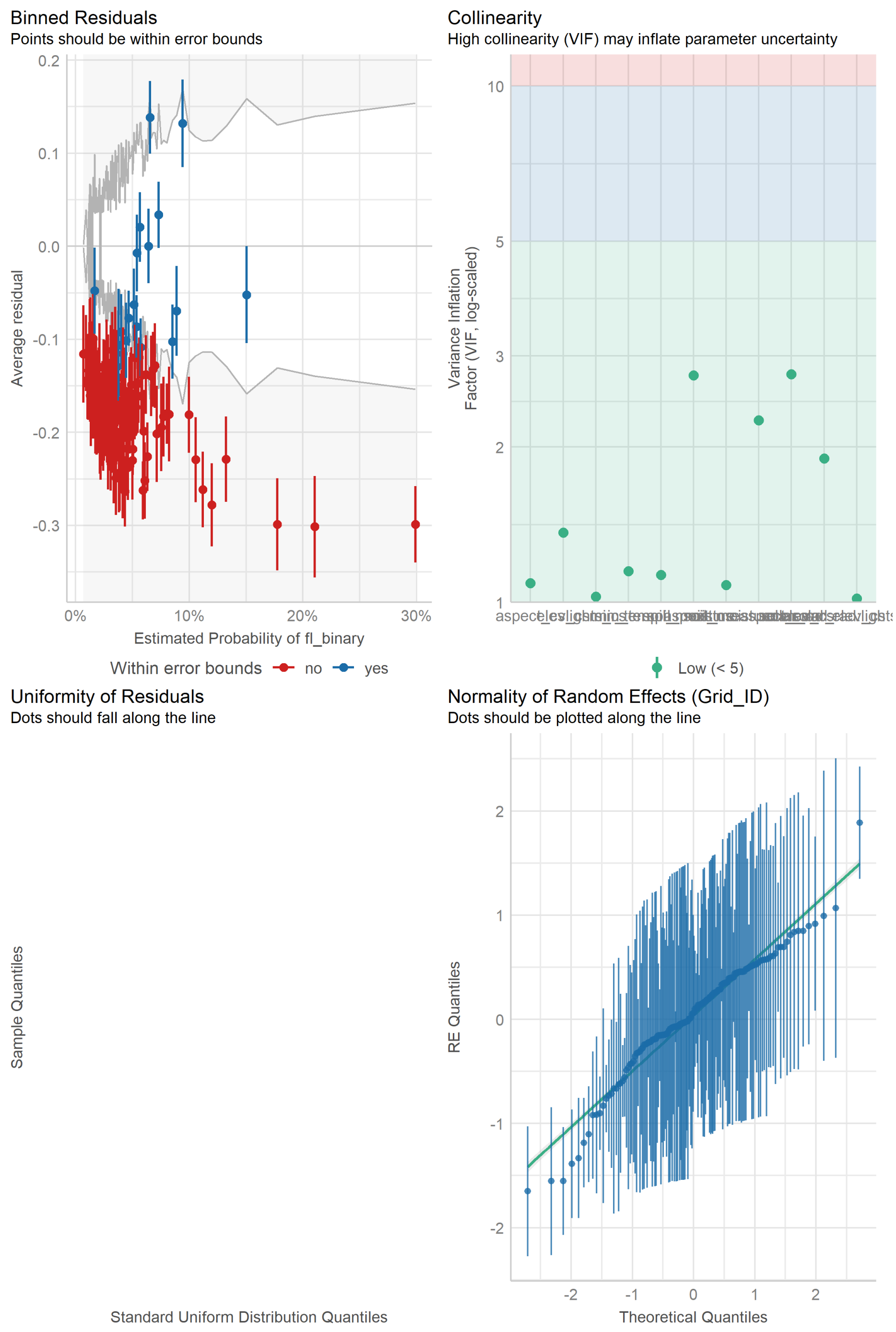

Figure 10. GLMM diagnostics for onset of flowering in mango

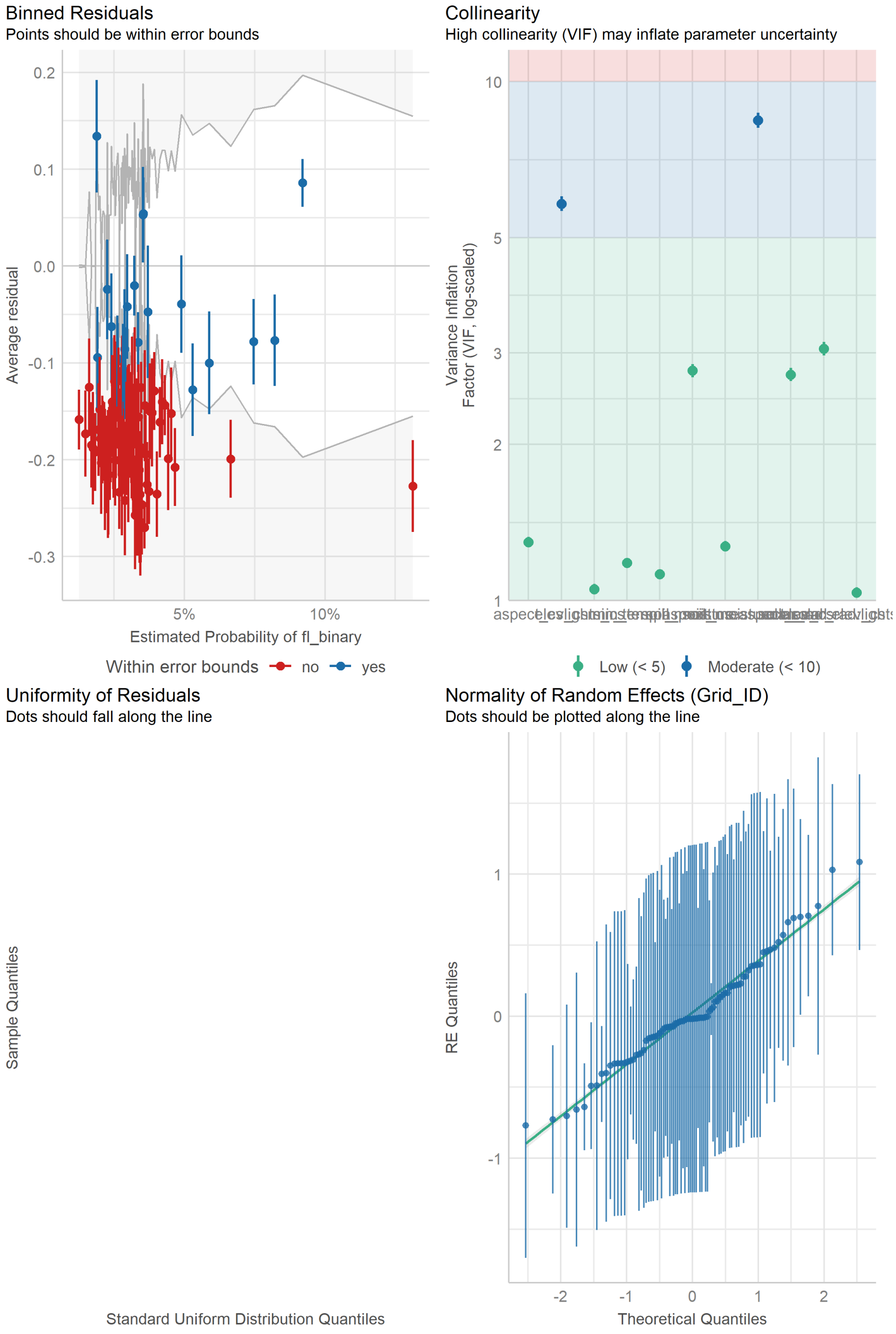

Figure S11. GLMM diagnostics for onset of flowering in tamarind

Table S8. Model coefficients for flowering onset in jackfruit with month as a predictor

### Fixed Effects

| Parameter | Log-Odds | SE | 95% CI | z | p |
|---|---|---|---|---|---|
| (Intercept) | -4.36 | 0.14 | (-4.64, -4.08) | -30.09 | < .001 |
| min temp | -0.70 | 0.05 | (-0.80, -0.60) | -13.87 | < .001 |
| soil moisture | -0.88 | 0.09 | (-1.05, -0.70) | -9.56 | < .001 |

| Parameter | Log-Odds | SE | 95% CI | z | p |
|---|---|---|---|---|---|
| solar rad | 0.05 | 0.06 | (-0.06, 0.16) | 0.93 | 0.350 |
| month | 0.17 | 0.01 | (0.15, 0.20) | 14.67 | < .001 |
| elev cs | -0.15 | 0.06 | (-0.27, -0.03) | -2.44 | 0.015 |
| aspect cs | -0.06 | 0.04 | (-0.15, 0.02) | -1.51 | 0.130 |
| lights cs | 0.03 | 0.06 | (-0.09, 0.15) | 0.47 | 0.635 |
| soil moisture × elev cs | -0.07 | 0.09 | (-0.24, 0.11) | -0.74 | 0.462 |
| solar rad × elev cs | -0.09 | 0.05 | (-0.18, 0.01) | -1.75 | 0.080 |
| min temp × aspect cs | -0.01 | 0.03 | (-0.07, 0.05) | -0.37 | 0.712 |
| soil moisture × aspect cs | -0.04 | 0.04 | (-0.11, 0.03) | -1.11 | 0.267 |
| solar rad × lights cs | 0.06 | 0.03 | (-2.92e-03, 0.13) | 1.87 | 0.061 |

### Random Effects

| Parameter | Coefficient |
|---|---|
| SD (Intercept: Grid_ID) | 0.86 |

Table S9. Model coefficients for flowering onset in mango with month as a predictor

### Fixed Effects

| Parameter | Log-Odds | SE | 95% CI | z | p |
|---|---|---|---|---|---|
| (Intercept) | -4.03 | 0.14 | (-4.29, -3.76) | -29.51 | < .001 |
| min temp | -0.78 | 0.05 | (-0.88, -0.67) | -14.63 | < .001 |
| soil moisture | -0.60 | 0.08 | (-0.76, -0.43) | -7.05 | < .001 |
| solar rad | 0.19 | 0.06 | (0.07, 0.30) | 3.23 | 0.001 |
| month | 0.13 | 0.01 | (0.11, 0.15) | 10.90 | < .001 |
| elev cs | -0.35 | 0.07 | (-0.50, -0.21) | -4.83 | < .001 |
| aspect cs | -0.02 | 0.04 | (-0.11, 0.07) | -0.42 | 0.674 |
| lights cs | 0.15 | 0.06 | (0.04, 0.26) | 2.62 | 0.009 |
| soil moisture × elev cs | 0.22 | 0.09 | (0.04, 0.39) | 2.38 | 0.017 |
| solar rad × elev cs | 0.07 | 0.06 | (-0.04, 0.18) | 1.18 | 0.239 |
| min temp × aspect cs | 0.01 | 0.04 | (-0.06, 0.08) | 0.30 | 0.763 |
| soil moisture × aspect cs | -0.03 | 0.04 | (-0.11, 0.04) | -0.89 | 0.375 |
| solar rad × lights cs | -0.06 | 0.03 | (-0.12, 5.70e-03) | -1.79 | 0.074 |

### Random Effects

| Parameter | Coefficient |
|---|---|
| SD (Intercept: Grid_ID) | 0.80 |

Table S10. Model coefficients for flowering onset in tamarind with month as a predictor

### Fixed Effects

| Parameter | Log-Odds | SE | 95% CI | z | p |
|---|---|---|---|---|---|
| (Intercept) | -3.13 | 0.17 | (-3.46, -2.80) | -18.77 | < .001 |
| min temp | 0.23 | 0.08 | (0.08, 0.39) | 2.93 | 0.003 |

| Parameter | Log-Odds | SE | 95% CI | z | p |
|---|---|---|---|---|---|
| soil moisture | 0.39 | 0.16 | (0.08, 0.69) | 2.49 | 0.013 |
| solar rad | 0.04 | 0.09 | (-0.14, 0.22) | 0.46 | 0.645 |
| month | -0.03 | 0.02 | (-0.07, 0.01) | -1.39 | 0.164 |
| elev cs | -0.06 | 0.16 | (-0.37, 0.26) | -0.35 | 0.729 |
| aspect cs | -0.07 | 0.07 | (-0.22, 0.07) | -0.96 | 0.335 |
| lights cs | -6.66e-04 | 0.07 | (-0.15, 0.14) | -8.97e-03 | 0.993 |
| soil moisture × elev cs | 0.12 | 0.17 | (-0.21, 0.45) | 0.73 | 0.468 |
| solar rad × elev cs | 0.07 | 0.08 | (-0.09, 0.23) | 0.88 | 0.376 |
| min temp × aspect cs | -0.03 | 0.06 | (-0.14, 0.08) | -0.55 | 0.585 |
| soil moisture × aspect cs | -0.04 | 0.07 | (-0.17, 0.09) | -0.59 | 0.556 |
| solar rad × lights cs | -3.24e-04 | 0.05 | (-0.09, 0.09) | -6.89e-03 | 0.994 |

### Random Effects

| Parameter | Coefficient |
|---|---|
| SD (Intercept: Grid_ID) | 0.64 |

#### Section S6. Leave-one-out cross validation for GLMMs

Table S11. Model coefficients for LOOCV models of flowering onset in jackfruit

| var | 2014 | 2015 | 2016 | 2017 | 2018 | 2019 | 2020 | 2021 | 202 |
|---|---|---|---|---|---|---|---|---|---|
| (Intercept) | -3.19 (0.11) | -3.16 (0.11) | -3.19 (0.12) | -3.14 (0.11) | -3.07 (0.11) | -3.19 (0.12) | -3.08 (0.11) | -3.08 (0.11) | -3.12 |
| min_temp | -0.65 (0.05) | -0.64 (0.05) | -0.68 (0.05) | -0.66 (0.05) | -0.63 (0.05) | -0.72 (0.06) | -0.59 (0.05) | -0.61 (0.05) | -0.65 |
| soil_moisture | -0.21 (0.08) | -0.26 (0.08) | -0.22 (0.08) | -0.25 (0.08) | -0.38 (0.09) | -0.29 (0.09) | -0.24 (0.08) | -0.19 (0.08) | -0.28 |
| solar_rad | 0.09 (0.06) | 0.08 (0.06) | 0.06 (0.06) | 0.08 (0.06) | 0 (0.06) | 0.05 (0.06) | 0.1 (0.06) | 0.18 (0.06) | 0.07 |
| elev_cs | -0.15 (0.06) | -0.17 (0.07) | -0.12 (0.06) | -0.15 (0.06) | -0.27 (0.09) | -0.19 (0.07) | -0.15 (0.06) | -0.19 (0.07) | -0.17 |
| aspect_cs | -0.06 (0.04) | -0.1 (0.05) | -0.08 (0.04) | -0.08 (0.04) | -0.05 (0.05) | -0.09 (0.05) | -0.05 (0.05) | -0.14 (0.05) | -0.07 |
| lights_cs | 0.06 (0.06) | 0.03 (0.06) | 0.03 (0.07) | 0.05 (0.06) | 0.06 (0.06) | 0.08 (0.07) | 0.12 (0.06) | 0.05 (0.07) | 0.05 |
| soil_moisture:elev_cs | -0.08 (0.09) | -0.06 (0.1) | -0.13 (0.09) | -0.09 (0.1) | 0.12 (0.11) | -0.07 (0.1) | 0 (0.09) | -0.03 (0.1) | -0.04 |
| solar_rad:elev_cs | -0.11 (0.05) | -0.08 (0.05) | -0.11 (0.05) | -0.13 (0.06) | -0.03 (0.05) | -0.12 (0.06) | -0.07 (0.05) | -0.13 (0.05) | -0.1 |
| min_temp:aspect_cs | -0.03 (0.03) | -0.03 (0.03) | -0.03 (0.03) | -0.04 (0.03) | -0.03 (0.03) | -0.02 (0.03) | -0.03 (0.03) | -0.05 (0.03) | -0.04 |
| soil_moisture:aspect_cs | -0.02 (0.04) | -0.01 (0.04) | -0.05 (0.04) | -0.02 (0.04) | 0 (0.04) | -0.04 (0.04) | -0.02 (0.04) | -0.07 (0.04) | -0.04 |
| solar_rad:lights_cs | 0.06 (0.03) | 0.09 (0.03) | 0.04 (0.03) | 0.08 (0.03) | 0.04 (0.03) | 0.07 (0.03) | 0.07 (0.03) | 0.02 (0.03) | 0.06 |

Table S12. Model coefficients for LOOCV models of flowering onset in mango

| var | 2014 | 2015 | 2016 | 2017 | 2018 | 2019 | 2020 | 2021 | 202 |
|---|---|---|---|---|---|---|---|---|---|
| (Intercept) | -3.15 (0.1) | -3.08 (0.1) | -3.22 (0.11) | -3.03 (0.1) | -3.07 (0.1) | -3.12 (0.1) | -3.08 (0.11) | -3.11 (0.11) | -3.05 |
| min_temp | -0.77 (0.05) | -0.73 (0.05) | -0.82 (0.06) | -0.71 (0.05) | -0.71 (0.05) | -0.79 (0.06) | -0.71 (0.05) | -0.71 (0.06) | -0.73 |
| soil_moisture | -0.16 (0.08) | -0.12 (0.08) | -0.05 (0.08) | -0.16 (0.08) | -0.31 (0.08) | -0.25 (0.08) | -0.29 (0.08) | -0.22 (0.08) | -0.24 |
| solar_rad | 0.21 (0.06) | 0.23 (0.06) | 0.22 (0.06) | 0.19 (0.06) | 0.13 (0.06) | 0.16 (0.06) | 0.15 (0.06) | 0.28 (0.07) | 0.18 |
| elev_cs | -0.34 (0.07) | -0.34 (0.07) | -0.44 (0.09) | -0.3 (0.07) | -0.34 (0.08) | -0.32 (0.07) | -0.29 (0.07) | -0.32 (0.1) | -0.35 |
| aspect_cs | -0.03 (0.04) | -0.04 (0.05) | -0.04 (0.05) | -0.04 (0.04) | -0.06 (0.05) | 0.01 (0.05) | -0.01 (0.05) | -0.05 (0.05) | -0.03 |
| lights_cs | 0.16 (0.06) | 0.18 (0.06) | 0.16 (0.06) | 0.16 (0.06) | 0.18 (0.06) | 0.15 (0.06) | 0.15 (0.06) | 0.18 (0.06) | 0.18 |
| soil_moisture:elev_cs | 0.18 (0.09) | 0.22 (0.1) | 0.3 (0.11) | 0.1 (0.1) | 0.25 (0.1) | 0.13 (0.1) | 0.13 (0.09) | 0.29 (0.1) | 0.21 |
| solar_rad:elev_cs | 0.05 (0.06) | 0.06 (0.06) | 0.06 (0.06) | 0.03 (0.06) | 0.06 (0.06) | 0.02 (0.06) | 0.03 (0.06) | 0.14 (0.07) | 0.06 |
| min_temp:aspect_cs | 0.01 (0.03) | 0.01 (0.04) | -0.01 (0.04) | -0.01 (0.04) | 0.04 (0.04) | 0.04 (0.04) | 0.04 (0.04) | 0.01 (0.04) | 0.01 |
| soil_moisture:aspect_cs | -0.01 (0.04) | 0.01 (0.04) | -0.04 (0.04) | -0.02 (0.04) | 0.01 (0.04) | -0.02 (0.04) | 0.01 (0.04) | -0.05 (0.04) | -0.01 |
| solar_rad:lights_cs | -0.06 (0.03) | -0.04 (0.03) | -0.08 (0.03) | -0.04 (0.03) | -0.06 (0.03) | -0.06 (0.03) | -0.05 (0.03) | -0.13 (0.04) | -0.04 |

Table S13. Model coefficients for LOOCV models of flowering onset in tamarind

| var | 2014 | 2015 | 2016 | 2017 | 2018 | 2019 | 2020 | 2021 | 202 |
|---|---|---|---|---|---|---|---|---|---|
| (Intercept) | -3.38 (0.14) | -3.37 (0.14) | -3.27 (0.14) | -3.29 (0.13) | -3.26 (0.14) | -3.31 (0.14) | -3.29 (0.14) | -3.15 (0.14) | -3.23 |
| min_temp | 0.18 (0.08) | 0.19 (0.08) | 0.24 (0.09) | 0.2 (0.08) | 0.24 (0.09) | 0.43 (0.1) | 0.25 (0.08) | 0.14 (0.09) | 0.2 ( |

| var | 2014 | 2015 | 2016 | 2017 | 2018 | 2019 | 2020 | 2021 | 202 |
|---|---|---|---|---|---|---|---|---|---|
| soil_moisture | 0.39 (0.15) | 0.29 (0.15) | 0.34 (0.15) | 0.25 (0.14) | 0.21 (0.15) | 0.45 (0.16) | 0.37 (0.15) | 0.18 (0.15) | 0.27 |
| solar_rad | 0.1 (0.09) | 0.1 (0.1) | 0.1 (0.09) | -0.02 (0.09) | 0.08 (0.1) | 0 (0.1) | 0.15 (0.1) | -0.08 (0.1) | 0.05 |
| elev_cs | -0.05 (0.16) | -0.13 (0.18) | 0.02 (0.15) | -0.13 (0.18) | -0.04 (0.17) | -0.01 (0.18) | -0.06 (0.17) | -0.03 (0.2) | -0.04 |
| aspect_cs | -0.09 (0.08) | -0.03 (0.08) | -0.13 (0.08) | -0.05 (0.08) | -0.09 (0.08) | -0.1 (0.08) | -0.1 (0.08) | -0.05 (0.08) | -0.06 |
| lights_cs | -0.02 (0.08) | -0.02 (0.08) | 0.02 (0.08) | 0 (0.07) | -0.03 (0.08) | 0.01 (0.08) | -0.04 (0.08) | 0.02 (0.08) | 0.02 |
| soil_moisture:elev_cs | 0.1 (0.17) | 0.21 (0.18) | 0.05 (0.17) | 0.18 (0.18) | 0.09 (0.18) | 0.14 (0.18) | 0.15 (0.17) | 0.02 (0.26) | 0.1 ( |
| solar_rad:elev_cs | 0.06 (0.08) | 0.1 (0.08) | 0.05 (0.08) | 0.09 (0.08) | 0.07 (0.09) | 0.08 (0.08) | 0.07 (0.08) | 0.03 (0.17) | 0.07 |
| min_temp:aspect_cs | -0.05 (0.06) | -0.02 (0.06) | -0.09 (0.06) | 0 (0.06) | -0.02 (0.06) | 0 (0.06) | -0.05 (0.06) | -0.05 (0.06) | -0.03 |
| soil_moisture:aspect_cs | -0.03 (0.07) | -0.07 (0.07) | -0.06 (0.07) | -0.07 (0.07) | -0.01 (0.07) | -0.01 (0.07) | -0.03 (0.07) | -0.04 (0.07) | -0.04 |
| solar_rad:lights_cs | -0.02 (0.05) | 0 (0.05) | 0 (0.05) | -0.01 (0.05) | 0.02 (0.05) | 0.05 (0.05) | -0.03 (0.05) | 0.01 (0.06) | 0.01 |

Table S14. Out-of-bag predictive power for LOOCV models of flowering onset in jackfruit

| year | cor | mse |
|---|---|---|
| 2014 | 0.2004693 | 0.0584499 |
| 2015 | 0.1217795 | 0.0432738 |
| 2016 | 0.1265467 | 0.0467056 |
| 2017 | 0.0935850 | 0.0384501 |
| 2018 | 0.1332462 | 0.0518880 |
| 2019 | 0.1155356 | 0.0457906 |
| 2020 | 0.0905950 | 0.0340696 |
| 2021 | 0.0539513 | 0.0320582 |
| 2022 | 0.1485688 | 0.0601856 |

Table S15. Out-of-bag predictive power for LOOCV models of flowering onset in mango

| year | cor | mse |
|---|---|---|
| 2014 | 0.1026427 | 0.0654172 |
| 2015 | 0.1699537 | 0.0388611 |
| 2016 | 0.0948727 | 0.0517563 |
| 2017 | 0.1589169 | 0.0341097 |
| 2018 | 0.1419168 | 0.0555414 |
| 2019 | 0.1498177 | 0.0356736 |
| 2020 | 0.1218867 | 0.0416410 |
| 2021 | 0.0684566 | 0.0334110 |
| 2022 | 0.2267429 | 0.0428476 |

Table S16. Out-of-bag predictive power for LOOCV models of flowering onset in tamarind

| year | cor | mse |
|---|---|---|
| 2014 | 0.0235131 | 0.0668460 |
| 2015 | 0.1197022 | 0.0668705 |
| 2016 | 0.0557611 | 0.0337775 |
| 2017 | 0.1026259 | 0.0303872 |
| 2018 | 0.0479079 | 0.0454108 |
| 2019 | 0.0025457 | 0.0307121 |
| 2020 | -0.0085910 | 0.0246159 |
| 2021 | 0.0165670 | 0.0262515 |
| 2022 | 0.0621319 | 0.0262394 |

#### Section S7. Supplementary methods and results for hotspot analysis

The emerging hotspot analysis from ArcGIS Pro was applied to the daily climate data (i.e., total precipitation, mean temperature, min temperature, max temperature, total solar radiation, and mean soil moisture) and the number of tree observations from 2014-2022. The emerging hotspot analysis evaluates spatiotemporal patterns in a variable using a combination of two statistical measures 1) the Getis-Ord Gi* statistic to identify hot spots and cold spots of climate measurements for each time step (i.e., fortnightly), and 2) the Mann Kendall trend test to examine how hot spots and cold spots have evolved over time.

The term 'hot spot' has been used generically across disciplines to describe a location has a value that is higher relative to its surroundings. However, a location with a high value may not be a statistically significant hot spot. For example, a hot spot of mean temperature was defined as an area that have a high value of mean temperature and is surrounded by other areas with high values as well.

##### Workflow for emerging hot spot analysis

Create space-time cube based on time-series daily climate/tree data at the pixel-level and choose 2 weeks as time step interval (i.e., aggregate daily data to the fortnightly level). The space-time cube stores space as latitude and longitude coordinates and time as another dimension. Each space-time bin in the output cube refers to a single cell for a single time interval (fortnightly).

Run emerging hotspot analysis based on aggregated fortnightly climate/tree variables from the space-time cube.

##### Category Definitions

**Last time step is hot:**

- New: the most recent time step interval is hot for the first time
- Consecutive: a single uninterrupted run of hot time step intervals, comprised of less than 90% of all intervals
- Intensifying: at least 90% of the time step intervals are hot, and becoming hotter over time
- Persistent: at least 90% of the time step intervals are hot, with no trend up or down
- Diminishing: at least 90% of the time step intervals are hot, and becoming less hot over time
- Sporadic: some of the time step intervals are hot
- Oscillating: some of the time step intervals are hot, some are cold

**Last time step is not hot:**

- Historical: at least 90% of the time step intervals are hot, but the most recent time step interval is not

**Last time step is cold:**

- New: the most recent time step interval is cold for the first time
- Consecutive: a single uninterrupted run of cold time step intervals, comprised of less than 90% of all
- Intensifying: at least 90% of the time step intervals are cold, and becoming colder over time
- Persistent: at least 90% of the time step intervals are cold, with no trend up or down
- Diminishing: at least 90% of the time step intervals are cold, and becoming less cold over time intervals
- Sporadic: some of the time step intervals are cold
- Oscillating: some of the time step intervals are cold, some are hot

**Last time step is not cold:**

- Historical: at least 90% of the time step intervals are cold, but the most recent time step interval is not

Table S17. Space-time cube characteristics from hotspot analysis of climate variables

| | |
|---|---|
| Number of time steps | 218 |
| Time step interval | 2 weeks |
| Time step alignment | End |

| | |
|---|---|
| Coordinate System | Asia South Albers Equal Area Conic |
| Cube extent across space | (coordinates in meters) |
| Min X | -5616423.582 |
| Min Y | 2211660.96 |
| Max X | -5275029.64 |
| Max Y | 2652552.121 |
| Locations | 285 |
| Total number | 7 |
| Total observations | 62130 |
| Total number | 1526 |

Overall Data Trend in MINTEMP

| | |
|---|---|
| Trend direction | Not Significant |
| Trend statistic | -0.2378 |
| Trend p-value | 0.812 |

Overall Data Trend in MEANSM

| | |
|---|---|
| Trend direction | Not Significant |
| Trend statistic | 0.2898 |
| Trend p-value | 0.772 |

Overall Data Trend in SOLARRAD

| | |
|---|---|
| Trend direction | Not Significant |
| Trend statistic | -0.3623 |
| Trend p-value | 0.7172 |

Overall Data Trend in TEMPORAL AGGREGATION COUNT

| | |
|---|---|
| Trend direction | Increasing |
| Trend statistic | 1.7162 |
| Trend p-value | 0.0861 |
